# Variation in fine scale recombination rate in temperature-evolved *Drosophila melanogaster* populations in response to selection

**DOI:** 10.1101/2022.04.28.489929

**Authors:** Ari Winbush, Nadia D. Singh

## Abstract

Meiotic recombination plays a critical evolutionary role in maintaining fitness in response to selective pressures due to changing environments. Variation in recombination rate has been observed amongst and between species and populations and within genomes across numerous taxa. Studies have demonstrated a link between changes in recombination rate and selection but the extent to which fine scale recombination rate varies between evolved populations during the evolutionary period in response to selection is under active research. Here we utilize a set of three temperature-evolved *Drosophila melanogaster* populations that were shown to have diverged in several phenotypes including recombination rate based on the temperature regime in which they evolved. Using whole genome sequencing data of these populations, we generated fine scale recombination maps of the three populations. We compare recombination rates and patterns among the three populations and show that they have diverged at fine scales but are conserved at broader scales. We further demonstrate a correlation between recombination rates and genomic variation in the three populations and observe variation in putative warm-spots between the populations with these enhanced areas and associated genes overlapping areas previously shown to have diverged in the three populations due to selection. These data support the existence of recombination modifiers in these populations which are subject to selection during evolutionary change.

## Introduction

Homologous recombination is a critical process in which genetic material is transferred between nucleic acid strands. During meiosis, this exchange occurs between homologous chromosomes and is essential for proper chromosomal segregation. In addition, recombination plays a critical evolutionary role primarily through the disruption of linkage disequilibrium, creation of new haplotypes, and increasing genetic variation allowing for modifications to fitness (Hill and Robertson, 1966; Begun and Aquadro, 1992; Kliman and Hey 1993; Kong et al., 2004; Baudet et al., 2013). Likewise, the tight control of recombination rate is important in maintaining fitness while preventing aneuploidy (Barton and Charlesworth, 1998; Ritz et al., 2017; Stapley et al., 2017).

Given this important role, recombination rate variation has been studied extensively and in numerous organisms. These studies have demonstrated almost universally the existence of recombination rate variation across genomes, within and among populations, as well as sexes and species at both fine and broad scales (For example: Nachman, 2002; Crawford et al., 2004; Cirulli et al., 2007; Mancera et al., 2008; Dumont et al., 2009; Singh et al., 2009; Stevison and Noor, 2010; Hinch et al., 2011; Smukowski and Noor, 2011; Chan et al., 2012; Comeron et al., 2012; Manzano-Winkler et al., 2013; Singh et al., 2013; Singhal et al., 2015; Smukowski Heil et. al., 2015; Hunter et al., 2016; Stapley et al., 2017; Dreissig et al., 2019; Shanfelter et al., 2019; Apuli et al., 2020; Hasan et al., 2020; Samuk et al., 2020). Noteworthy among studies of recombination rate are 1) the conservation of intragenomic recombination rate variation and landscapes at broad scales between closely related species and populations and 2) the rapid divergence and evolution of recombination rate variation at fine scales. Examples of the former include the variation in recombination rate observed around heterochromatic chromosomal structures and genomic motifs; for example, suppressed recombination rate around centromeres compared to other regions in fruit flies and yeast or changes in recombination rate as a function of distance from telomeres in humans (Beadle, 1932; Nachman 2002; Anderson et al., 2003; Chen et al., 2008; Myers et al., 2008; Comeron et al., 2012). Concerning the latter, the existence of localized recombination “hotspots” consisting small regions in which recombination rate is multi-fold higher than background, has been observed in a variety of organisms. These hotspots appear to be relatively stable within populations yet vary significantly among species and populations which suggests a rapid evolution of divergence at this scale (Myers et al., 2005; Ptak et al., 2005; Winckler et al., 2005; Mancera et al., 2008; Baudat et al., 2010; Smagulova et al., 2011 Baudat et al., 2013).

Another trend that has emerged from decades of work on recombination rate is that populations subjected to selection on a particular phenotype have been shown to also exhibit differences in recombination. This is seen in the context both experimental evolution experiments and domestication. Concerning strong artificial selection due to domestication, studies have observed changes recombination rates, and polymorphisms in recombination-specific genes in domesticated plants and animals (Burt and Bell, 1987; Charlesworth, 1993; Ross-Ibarra, 2004; Groenen et al., 2009; Poissant et al., 2010; Sidhu et al., 2017; Dreissig et al., 2019). Nevertheless, this phenomenon is not always observed in studies of domestication and various underlying causes have been reported (Otto and Barton, 2001; Muñoz-Fuentes et al., 2014). In the contexts of both experimental evolution and studies of natural populations that have adapted to local environments and parasite-host coevolution, changes in recombination rate are also observed (Korol and Iliadi, 1994; Korol, 1999; Bourguet et al., 2003; Kerstes et al., 2012; Aggarwal et al., 2015; Aggarwal et al., 2019; Neupane and Xu, 2020).

Much study has therefore been devoted to elucidating the genetic basis and evolution of fine-scale recombination rate variation. An important discovery in this area was the histone methyltransferase-encoding gene, *PRDM9*, shown in humans, mice, and other taxa to regulate the location of recombination hotspots (Oliver et al. 2009; Baudat et al., 2010; Berg et al., 2011; Hinch et al. 2011; Schwartz et al., 2014). However, many organisms lack *PRDM9*, and evidence would suggest additional genetic determinants are responsible for fine scale recombination rate variation in these specimens.

*Drosophila* represents a commonly used model system for study of genetic determinants and evolution of fine scale recombination rate variation. Lacking *PRDM9* and hotspots at the same scale as observed in humans and mice, numerous studies nevertheless have demonstrated recombination rate variation at fine scales among *Drosophila* species and within genomes (Cirulli et al., 2007; Singh et al., 2009; Stevison and Noor, 2010; Chan et al., 2012; Comeron et al., 2012; Manzano-Winkler et al., 2013; Singh et al., 2013; Smukowski Heil et al., 2015; Adrion et al., 2020). There is likewise evidence to suggest the existence of recombination modifiers and that selection acts on recombination rate both directly and in the context of indirect selection of other traits in response to novel environments, experimental evolution, and artificial selection (Kidwell, 1972; Charlesworth, 1993; Korol and Iliadi, 1994; Bourget et al., 2003; Aggarwal et al., 2015; Smukowski Heil et al., 2015). However, a comprehensive analysis of recombination rate variation across and within species, populations and genomes, and factors driving fine scale recombination rate divergence and modifiers remains lacking in *Drosophila*. For example, the existence of recombination modifiers would suggest that under conditions promoting rapid evolution and selection, areas of enhanced recombination would overlap genomic regions regions undergoing selection but the extent to which this occurs is unknown. The examination of fine scale recombination rates in populations subject to selection during an experimental evolution period would provide insight into these questions.

We previously utilized a set of experimentally evolved *Drosophila melanogaster* populations that were subjected to one of three temperature treatments (Warm, Cold, and Temporally fluctuating (Temp)) over a three-year period and were subsequently demonstrated to show fixed heritable differences in recombination rate (Kohl and Singh, 2018). Those from the Warm and Cold regimes exhibited the highest and lowest rates respectively and the Temp lines exhibited an intermediate rate although no variation in recombination rate plasticity was observed (Kohl and Singh, 2018). Whole genome sequencing of these populations and subsequent analysis of single nucleotide polymorphism (SNP)-allele frequencies identified multiple regions of divergence between the three populations stemming from pairwise comparisons (Warm vs Cold, Cold vs Temp, and Warm Vs Temp) (Winbush and Singh, 2021). These regions of divergence overlapped regions of reduced nucleotide diversity and reduced empirical recombination rates suggesting that these regions were subject to selection during the experimental evolution period. Additionally, we were able to map divergent SNPs to potential candidate loci responsible for variation in recombination rate between the three populations (Winbush and Singh 2021). Interestingly, these loci showed overlap with those, previously identified in a separate screen of 205 inbred lines associated with the *D. melanogaster* Genetic Reference Panel (DGRP) investigating variation in recombination rate at the population level and suggested that candidate genes regulating recombination rate between populations and within populations might be shared (Hunter et al., 2016; Winbush and Singh 2021). Therefore, these three temperature-regime populations represent ideal candidates to test for both the divergence of recombination rate at fine scales in response to selection and potential overlap of localized areas of enhanced recombination rate with areas undergoing selection during the evolutionary period.

We took advantage of the existing linkage disequilibrium (LD)-based statistical software LD-helmet to infer historical patterns of fine scale recombination rate in these three populations. This offers advantages over empirical methods of assessing fine scale recombination rates using pedigree analyses which are often laborious requiring numerous controlled crosses and extensive genotyping. However, LD based methods also have limitations due to other factors that influence patterns of LD across the genome such as sudden demographic changes, genetic drift, selection, and mutation. Like previous studies we therefore compare our data to previously generated empirical data and show good correlation between the two. This result is consistent with other studies comparing LDhelmet-generated data to empirical results (Singh et al., 2009; Chan et al., 2012; Smukowski Heil et al., 2015) showing strong correlation and demonstrating the efficacy of this LD-based method (see results).

Our data show that recombination rates in these three populations have diverged in fine scales but are conserved at broader scales. We also demonstrate an increase in recombination rate in areas undergoing selection despite conservation of intragenomic recombination rates at broad scales. The overlap between areas of enhanced recombination and genomic regions subject to selection likewise supports the existence of recombination modifiers which are subject to selection during rapid evolutionary change.

## Materials and Methods

### Fly Populations

The *D. melanogaster* populations used in this study were generated previously as part of an experimental evolution study in which wild caught females from British Columbia were used to establish a large set of isofemale lines. From this large breeding population, three sets of five replicate populations were generated and each set was subjected over an approximately three-year period to one of three experimental temperature regimes: Warm (25°C), Cold (16°C) and Temporally Fluctuating (Temp; migration between the Warm and Cold regimes every four weeks) (Yeaman et al., 2010). At the end of this period, a set of isofemale lines were established from each population replicate and maintained under constant conditions over a 27-month period to allow for establishment of isogenic lines (Cooper et al., 2012). These resulting populations were previously shown to exhibit heritable differences in recombination rate with those from the Warm and Cold regimes exhibiting the highest and lowest rates respectively and the Temp regimes lines exhibiting an intermediate rate (Kohl and Singh, 2018).

### Crosses and Variant Calling

Generation, sequencing, and variant calling of haploid embryos was performed previously and is only described briefly here (Winbush and Singh, 2021). Females from isofemale lines representing the three sets of five replicate populations for each temperature regime were crossed to males bearing the male-sterile *ms(3)K81* mutation. Resulting progeny from this cross are haploid with all genetic material being maternally derived (Fuyama, 1984, Langley et al. 2011). Haploid embryos surviving to the first instar larval stage (approximately 1% of progeny) were collected from each of isofemale line resulting in 48 individual embryos for each temperature regime (hereafter referred to the Cold, Warm, and Temp populations).

Preparation, sequencing, and final coverage of DNA libraries from individual embryos was as described previously (Winbush and Singh, 2021). Resulting reads were mapped to the *D. melanogaster* reference genome (Flybase, r6.22) using the MosaikAligner toolset (v2.2.3; Lee et al., 2014). After removing samples with poor alignment, a total of 137 samples representing each embryo remained for downstream variant calling. Single-nucleotide polymorphism (SNP) calling was performed using both Freebayes (v.1.2.0) and Joint Genotyper for Inbred Lines (JGIL; v.1.6) software (Marth et al., 1999; Garrison and Marth, 2012; Stone, 2012). SNPs called by both methods were filtered as detailed previously which resulted in 850,298 SNPs in our dataset across the five major chromosome arms that were used in the previous study (Winbush and Singh, 2021). For this study we filtered an additional 15,091 variants that exhibited differences in alternate alleles called by Freebayes or JGIL resulting in 835,207 SNPs in our dataset.

### Generation of recombination estimate maps

Recombination rate estimates for the three populations were performed using the LDhelmet (v.1.10) program. Like LDhat but tailored to *Drosophila*, LDhelmet utilizes a reversible jump Markov Chain Monte Carlo (rjMCMC) model to estimate the population-scaled recombination rate, ρ, based on patterns of linkage disequilibrium (LD) among SNPs within a set of haplotypes comprising a population. Values of ρ (4N_e_r; N_e_=effective population size; r=recombination rate) are partially based on those required to produce observed rates of LD decay between loci within a population (Chan et al., 2012; Li and Stephens, 2003; Auton and McVean, 2007; McVean et al., 2004). Additionally, LDhelmet is suited to handling larger sample sizes, missing alleles, and the higher SNP densities common in *Drosophila* (Chan et al., 2012).

To generate FASTA files for input into LDhelmet, we used a combination of shell tools and bcftools (v.1.9-67-g626e46b; Li, 2011) to separate our initial VCF file by population (Cold, Warm, and Temp) and chromosome arm. For each population, a FASTA consensus sequence containing the corresponding SNPs, at the appropriate locations was created for each sample. This was performed separately for all five long chromosome arms resulting in a set of haplotypes for each population and chromosome arm. The use of haploid embryos allowed for a single haplotype per sample with no haplotype phasing requirements. As a result, complete population specific FASTA files for each chromosome contained 42, 47, and 48 individual sample haplotypes for the respective Cold, Warm and Temp populations.

We ran LDhelmet individually for each chromosome arm and population using the default parameters for the find_conf and pade modules. Theta values were estimated using the PopGenome package implemented in R (v.2.7.5; Pfeifer et al., 2014) with values at 0.001 being observed for the three populations. For the mutation transition matrix, we used the values previously derived from the Raleigh population in Chan et al. (2012) as we expect our populations to have similar values based on the similar outgroup reference genomes. We ran the rjmcmc module for 1,000,000 iterations with a 100,000-iteration burn-in. The choice of block penalty has a large impact on recombination estimate map smoothing. Previous studies in *Drosophila* species utilized a block penalty of 50 and found no effect of different block penalties on overall results (Smukowski Heil et al., 2015; Chan et al., 2012). We therefore used a block penalty of 50.

### Empirical Recombination maps

We utilized empirical fine scale recombination maps that were previously generated for *D. melanogaster* through large scale crosses and sequencing of a set of inbred strains (Comeron et al., 2012). Recombination maps from this data were generated at 100kb intervals. Comparison to our LD based data at similar intervals was performed by first converting our population scaled rates (ρ/bp) to cM/Mb using methods previously described (Chan et al., 2012; Smukowski Heil et al., 2015), and averaging over the same interval.

### Comparison of recombination rates between populations

Recombination rates between populations were compared at the chromosome level at different scales using average rates over non-overlapping intervals. Population comparisons (Cold vs Warm, Cold vs Temp, Warm vs Temp) were performed using Spearman’s rank coefficient and are reported as Spearman’s rho and P-value at the indicated interval size. We also assessed variation of all intervals among populations using ANOVA and report individual comparisons between populations using Tukey’s HSD test for each pairwise comparison.

## Results

### Comparison to Empirical Recombination Rates

We utilized the LDhelmet software to infer recombination rate histories within our temperature-derived experimental evolution populations of which the resulting fine-scale recombination maps are shown in Figure 1. These populations were previously shown to have diverged in several phenotypes in addition to recombination rate including cell membrane lipid composition, cell size, metabolism, fecundity, developmental plasticity, and thermal tolerance (Cooper et al., 2012; Condon et al., 2014; Condon et al., 2015; Adrian et al., 2016; Alton et al., 2017; Le Vinh Thuy et al., 2016, Kohl and Singh, 2018). They therefore present an excellent opportunity to utilize LD-based methodologies to infer historical recombination rates within the different populations. We also note that these estimates of recombination will be based on the shared evolutionary history of these populations, given they were founded from the same source, in addition to changes in recombination during the course of the experimental evolution study. LD-based methods of assessing recombination rate offers practical advantages over empirical methods such as controlled crosses and extensive genotyping which are often laborious. However, as noted in Smukowski Heil et al. 2015, statistical LD based methods have their limitations due other factors that influence LD including sudden demographic changes, genetic drift, changing mutation rates, and selection (Smith and Fearnhead, 2005; Slatkin, 2008; Dapper and Payseur, 2018; van Eeden et al., 2021). Furthermore, LD-based methods are typically utilized to infer historical recombination rates over long timespans into the past (Hermann et al., 2019; van Eeden et al., 2021) which contrasts the relatively short experimental-evolution time frame of our populations.

**Figure 1.**
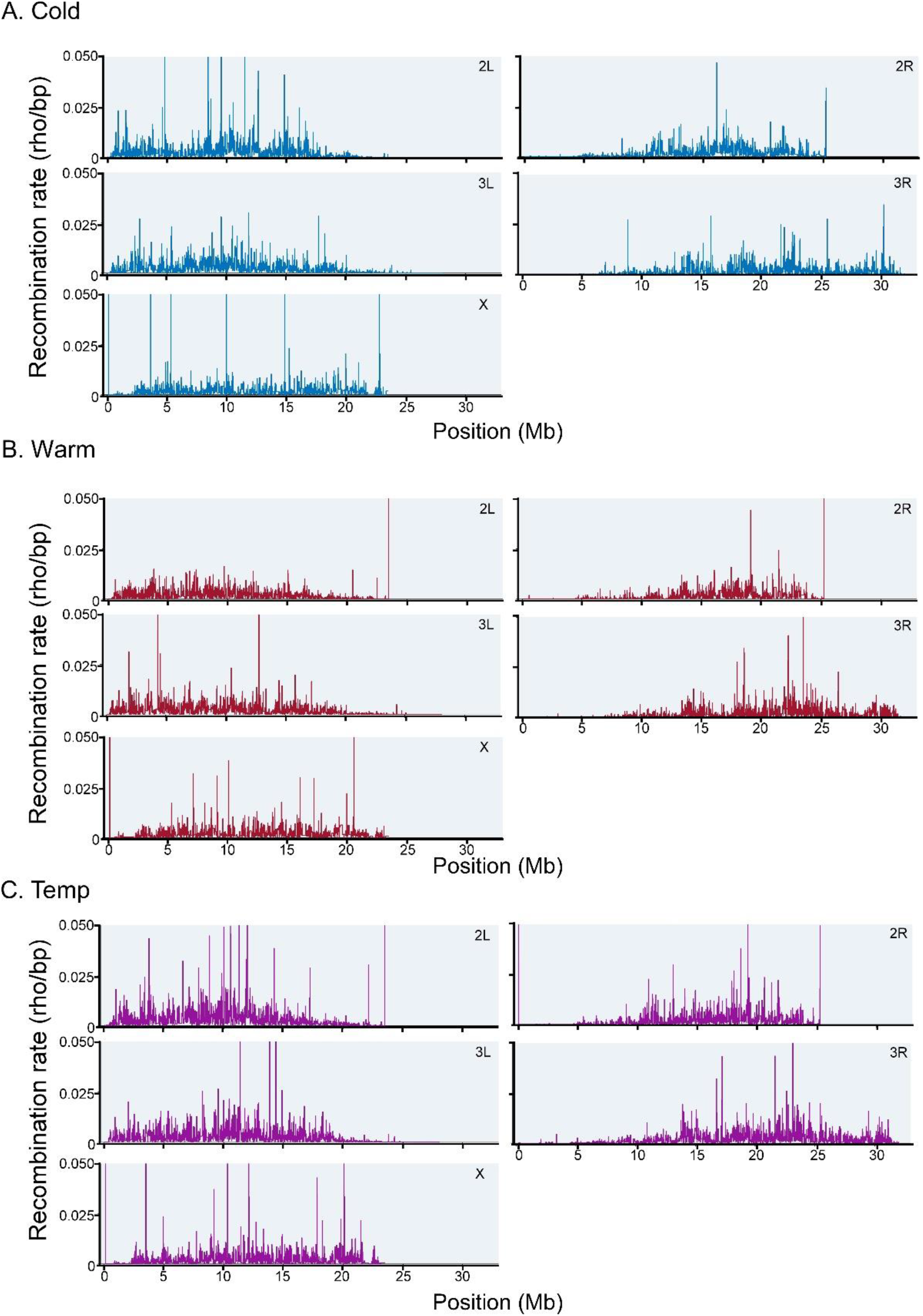
LDhelmet’s fine scale recombination estimate maps. (A) Cold population. (B) Warm population. (C) Temp population.

We therefore compared our LDhelmet data for the three populations to recombination maps produced based on publicly available empirical data (Comeron et al. 2012; Fiston-Lavier et al., 2010). Comparisons based on average recombination rates at 100kb and 200kb intervals showed significant correlation at both scales (Table 1). We note stronger correlation values (Spearman’s, rho) at the broader scale of 200kb for all three populations and all five major chromosome arms (Table 1B). In all three populations, the highest correlations were found overall on chromosome arms 2L and 3L while chromosome X had the weakest correlations. In comparison to results obtained by the Chan et al. 2012 study which utilized a different empirical dataset, and populations derived from Raleigh, USA (RA) and Gikongoro, Rwanda (RG), we note slightly weaker correlations overall of our three populations to the empirical dataset we utilized. For example, mean correlations at 200kb obtained by Chan et al. for RA and RG were 0.74, 0.77, 0.64, 0.69 and 0.67 for chromosomes 2L, 2R, 3L, 3R, and X respectively, while mean values for the Cold Warm and Temp populations were 0.71, 0.50, 0.72, 0.61, 0.43 for the same respective chromosomes. The use of different empirical datasets and different populations accounts for some of these differences; however, we should note that our experimental evolution design, which made use of different rearing temperatures likely affects recombination rates and is therefore likely also responsible. For example, we previously noted only a small overlap in SNPs in our populations to those of the *D. melanogaster* Genetic Research Panel (DGRP) which are also derived from the RA population.

**Table 1.** Comparison of LDhelmet-estimated recombination rates and empirical data derived from Comeron et al. (2012) for the three populations (Cold, Warm, Temp) for the five major chromosomes at the (A) 100kb interval and (B) 200kb interval. Entries (separated by semicolons) as follows: Spearman’s rho, P-value, and average number of SNPs per interval.

|  | Cold | Warm | Temp |
| --- | --- | --- | --- |
| **2L** | 0.61, P<0.0001 | 0.66, P<0.0001 | 0.63, P<0.0001 |
| **2R** | 0.50, P<0.0001 | 0.48, P<0.0001, | 0.48, P<0.0001 |
| **3L** | 0.57, P<0.0001 | 0.61, P<0.0001 | 0.65, P<0.0001 |
| **3R** | 0.55, P<0.0001 | 0.54, P<0.0001 | 0.53, P<0.0001 |
| **X** | 0.32, P<0.0001 | 0.38, P<0.0001 | 0.42, P<0.0001 |

Table 1. Comparison of LDhelmet-estimated recombination rates and empirical data derived from Comeron et al.
|  | Cold | Warm | Temp |
| --- | --- | --- | --- |
| **2L** | 0.72, P<0.0001 | 0.72, P<0.0001 | 0.70, P<0.0001 |
| **2R** | 0.52, P<0.0001 | 0.47, P<0.0001 | 0.52, P<0.0001 |
| **3L** | 0.70, P<0.0001 | 0.72, P<0.0001 | 0.75, P<0.0001 |
| **3R** | 0.62, P<0.0001 | 0.61, P<0.0001 | 0.60, P<0.0001 |
| **X** | 0.38, P<0.0001 | 0.44, P<0.0001 | 0.46, P<0.0001 |

We note several features across our datasets that are similar to those found in the empirical dataset. For example, we observed decreased recombination rates toward centromeric regions in our three populations as has been noted in other studies (Comeron et al., 2012; reviewed in Nachman, 2002) and a notable spike in recombination on chromosome 2L near the 10 Mb position (Chan et al., 2012) which we note also occurred in the Cold and Temp populations but not the Warm population (Figure 2). Uniform variation in recombination rate is noted on the X chromosome in all three populations but the large-scale fluctuations noted in the empirical dataset and previously in the flybase dataset (Chan et al., 2012) are less prominent. Mean recombination rates for all 200kb intervals on the X chromosome were also lower for the three populations overall (2.89 cM/Mb, 2.88 cM/Mb and 2.88 cM/Mb for the Cold, Warm and Temp populations respectively vs 2.95 cM/Mb for the empirical dataset). However, we also note two large spikes in rates near the start of the X chromosome in both the Warm and Temp populations and a notable spike on chromosome arm 2L and X in the Cold population (Figure 1). In summary, our data suggests a good estimation of recombination rates in our three populations with some expected variation which is further explored below.

**Figure 2.**
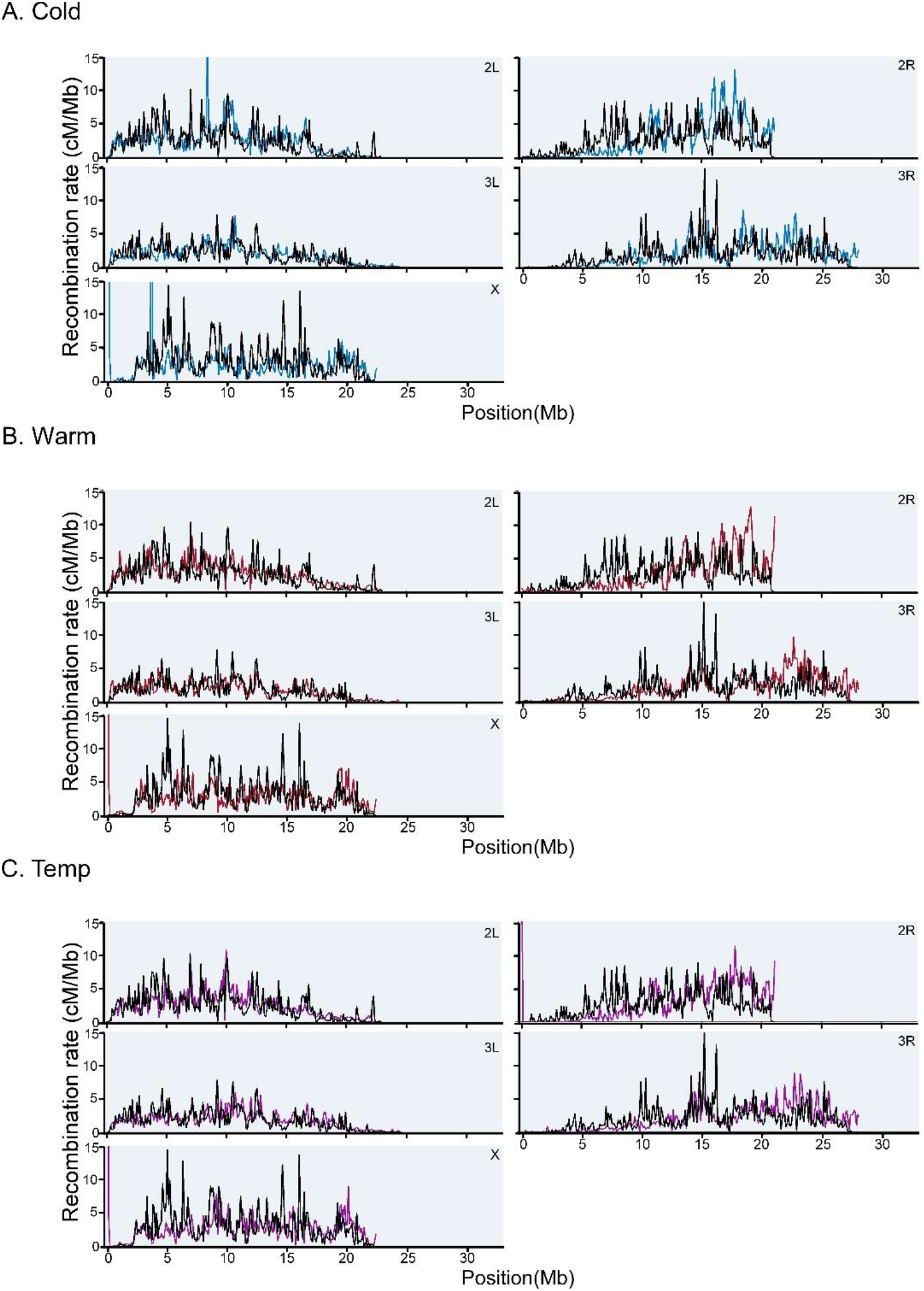
A comparison of LDhelmet estimates of recombination rate to those derived from empirical methods along the five major chromosome arms. Empirical data (black lines) are derived from Comeron et al., (2012) at 100kb scales with LDhelmet estimates at the same scale. (A) Cold population LDhelmet estimate (blue line) and empirical data. (B) Warm population LDhelmet estimate (red line) and empirical data. (C) Temp population LDhelmet estimate (purple) and empirical data.

### Recombination Rate Variation between Populations

Based on the experimental evolution design that generated these populations we expect to find variation in historical recombination rate patterns as well as more recent effects of adaption to temperature on recombination rate. Previous studies found variation in recombination rate at fine scales between different *Drosophila* species and different *D. melanogaster* populations (Chan et al., 2012; Smukowski Heil et al. 2015; Adrion et al., 2020). We therefore tested our populations for variation in recombination rate at fine and relatively broad scales. Intervals utilized and average number of SNPs per interval are summarized in Table 2. Examining average rates at 50kb intervals (Figure 3) we note that all three populations are statistically correlated overall in each of the comparisons (Cold vs Warm, Cold vs Temp, Warm vs Temp; Table 3A). However, consistently lower correlations were observed across all five chromosomes for the comparisons that include the Cold population (Cold vs Warm, Cold vs Temp). In contrast, the Warm vs Temp comparison showed the highest correlation coefficients across all five chromosomes suggesting greater variation in recombination rate within the Cold population. Interestingly, variation in recombination rate is also observed between chromosomes when populations are compared. This is most noticeable on the X chromosome which consistently had the lowest correlation values across all three comparisons (Table 3A). Significant differences in mean recombination rates are also observed across the three populations for chromosome arms 2L, 3L, and 3R (P<.0001; one-way analysis of variance). Post hoc comparisons of population pairs confirm these results and would suggest higher recombination rates in the Temp population (Figure 4).

**Table 2.** Average number of SNPs per interval for a given interval size for the three populations and chromosome arms.

|  | Cold | Warm | Temp |
| --- | --- | --- | --- |
| **2L** | 50kb: 306<br>100 kb: 613<br>200kb: 1227<br>500kb: 3111 | 50kb: 337<br>100 kb: 675<br>200kb: 1351<br>500kb: 3427 | 50kb: 361<br>100 kb: 724<br>200kb: 1449<br>500kb: 3676 |
| **2R** | 50kb: 200<br>100 kb: 400<br>200kb: 800<br>500kb: 1989 | 50kb: 204<br>100 kb: 408<br>200kb: 816<br>500kb: 2031 | 50kb: 225<br>100 kb: 450<br>200kb: 901<br>500kb: 2240 |
| **3L** | 50kb: 279<br>100 kb: 559<br>200kb: 1122<br>500kb: 2805 | 50kb: 290<br>100 kb: 581<br>200kb: 1167<br>500kb: 2918 | 50kb: 311<br>100 kb: 623<br>200kb: 1251<br>500kb: 3129 |
| **3R** | 50kb: 227<br>100 kb: 455<br>200kb: 910<br>500kb: 2275 | 50kb: 234<br>100 kb: 468<br>200kb: 937<br>500kb: 2344 | 50kb: 251<br>100 kb: 503<br>200kb: 1007<br>500kb: 2518 |
| **X** | 50kb: 192<br>100 kb: 385<br>200kb: 775<br>500kb: 1929 | 50kb: 201<br>100 kb: 402<br>200kb: 809<br>500kb: 2014 | 50kb: 215<br>100 kb: 430<br>200kb: 864<br>500kb: 2151 |

**Table 3.**
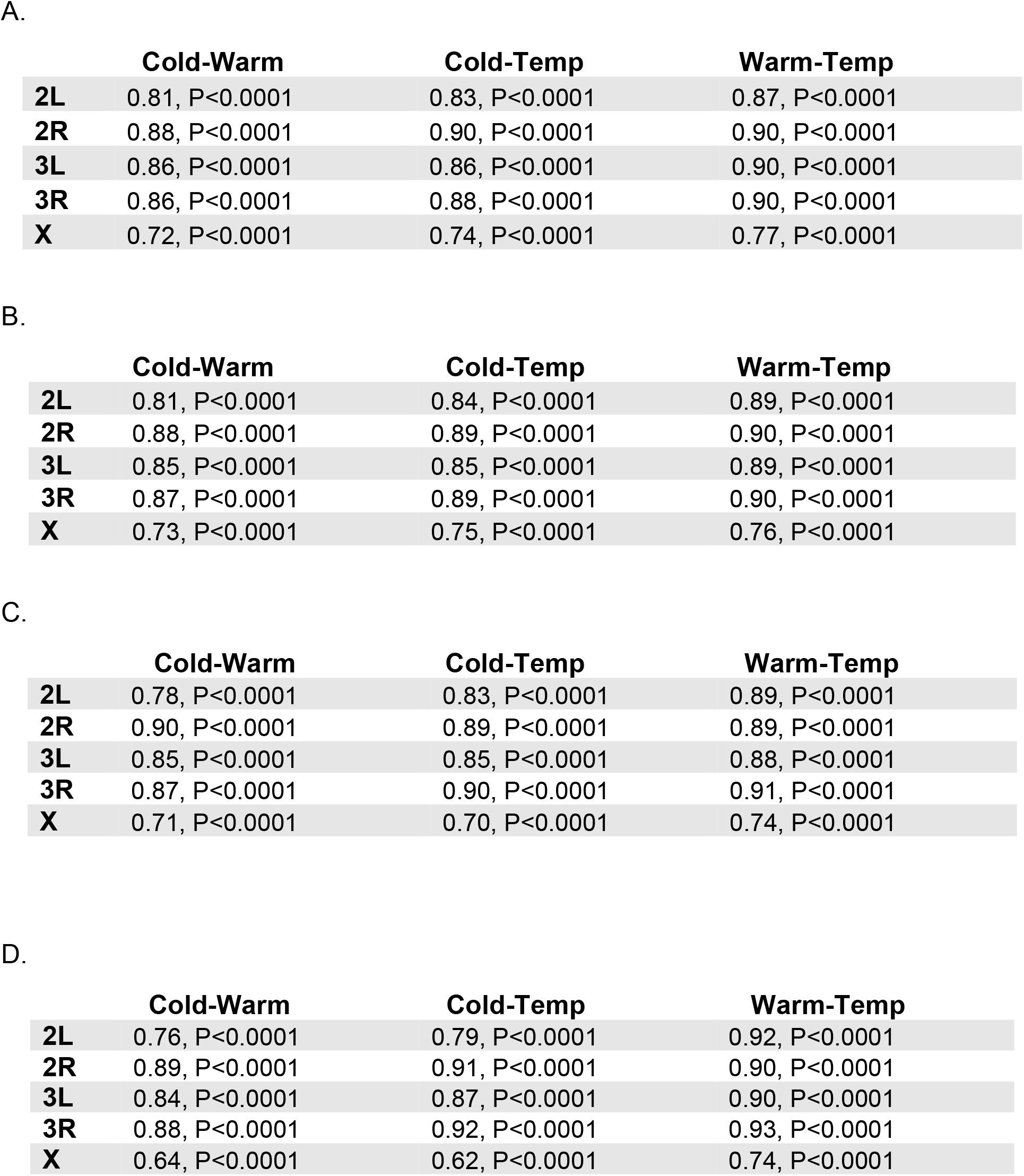
Comparison of LDhelmet-estimated recombination rates for the three population comparisons (Cold vs Warm, Cold vs Temp, Warm vs Temp) for the five major chromosome arms at (A) 50kb interval, (B) 100kb interval, (C) 200kb, and (D) 500kb interval. Entries in first row for each chromosome arm (separated by semicolons) as follows: Spearman’s rho, P-value while second row entries (separated by semicolons) give average number of SNPs per interval for the appropriate population.

|  | Cold-Warm | Cold-Temp | Warm-Temp |
| --- | --- | --- | --- |
| **2L** | 0.81, P<0.0001 | 0.83, P<0.0001 | 0.87, P<0.0001 |
| **2R** | 0.88, P<0.0001 | 0.90, P<0.0001 | 0.90, P<0.0001 |
| **3L** | 0.86, P<0.0001 | 0.86, P<0.0001 | 0.90, P<0.0001 |
| **3R** | 0.86, P<0.0001 | 0.88, P<0.0001 | 0.90, P<0.0001 |
| **X** | 0.72, P<0.0001 | 0.74, P<0.0001 | 0.77, P<0.0001 |

B.
|  | Cold-Warm | Cold-Temp | Warm-Temp |
| --- | --- | --- | --- |
| **2L** | 0.81, P<0.0001 | 0.84, P<0.0001 | 0.89, P<0.0001 |
| **2R** | 0.88, P<0.0001 | 0.89, P<0.0001 | 0.90, P<0.0001 |
| **3L** | 0.85, P<0.0001 | 0.85, P<0.0001 | 0.89, P<0.0001 |
| **3R** | 0.87, P<0.0001 | 0.89, P<0.0001 | 0.90, P<0.0001 |
| **X** | 0.73, P<0.0001 | 0.75, P<0.0001 | 0.76, P<0.0001 |

C.
|  | Cold-Warm | Cold-Temp | Warm-Temp |
| --- | --- | --- | --- |
| **2L** | 0.78, P<0.0001 | 0.83, P<0.0001 | 0.89, P<0.0001 |
| **2R** | 0.90, P<0.0001 | 0.89, P<0.0001 | 0.89, P<0.0001 |
| **3L** | 0.85, P<0.0001 | 0.85, P<0.0001 | 0.88, P<0.0001 |
| **3R** | 0.87, P<0.0001 | 0.90, P<0.0001 | 0.91, P<0.0001 |
| **X** | 0.71, P<0.0001 | 0.70, P<0.0001 | 0.74, P<0.0001 |

Table 3. Comparison of LDhelmet-estimated recombination rates for the three population comparisons (Cold vs Warm, Cold vs Temp, Warm vs Temp) for the five major chromosome arms at (A) 50kb interval, (B) 100kb interval, (C) 200kb, and (D) 500kb interval.
|  | Cold-Warm | Cold-Temp | Warm-Temp |
| --- | --- | --- | --- |
| **2L** | 0.76, P<0.0001 | 0.79, P<0.0001 | 0.92, P<0.0001 |
| **2R** | 0.89, P<0.0001 | 0.91, P<0.0001 | 0.90, P<0.0001 |
| **3L** | 0.84, P<0.0001 | 0.87, P<0.0001 | 0.90, P<0.0001 |
| **3R** | 0.88, P<0.0001 | 0.92, P<0.0001 | 0.93, P<0.0001 |
| **X** | 0.64, P<0.0001 | 0.62, P<0.0001 | 0.74, P<0.0001 |

**Figure 3.**
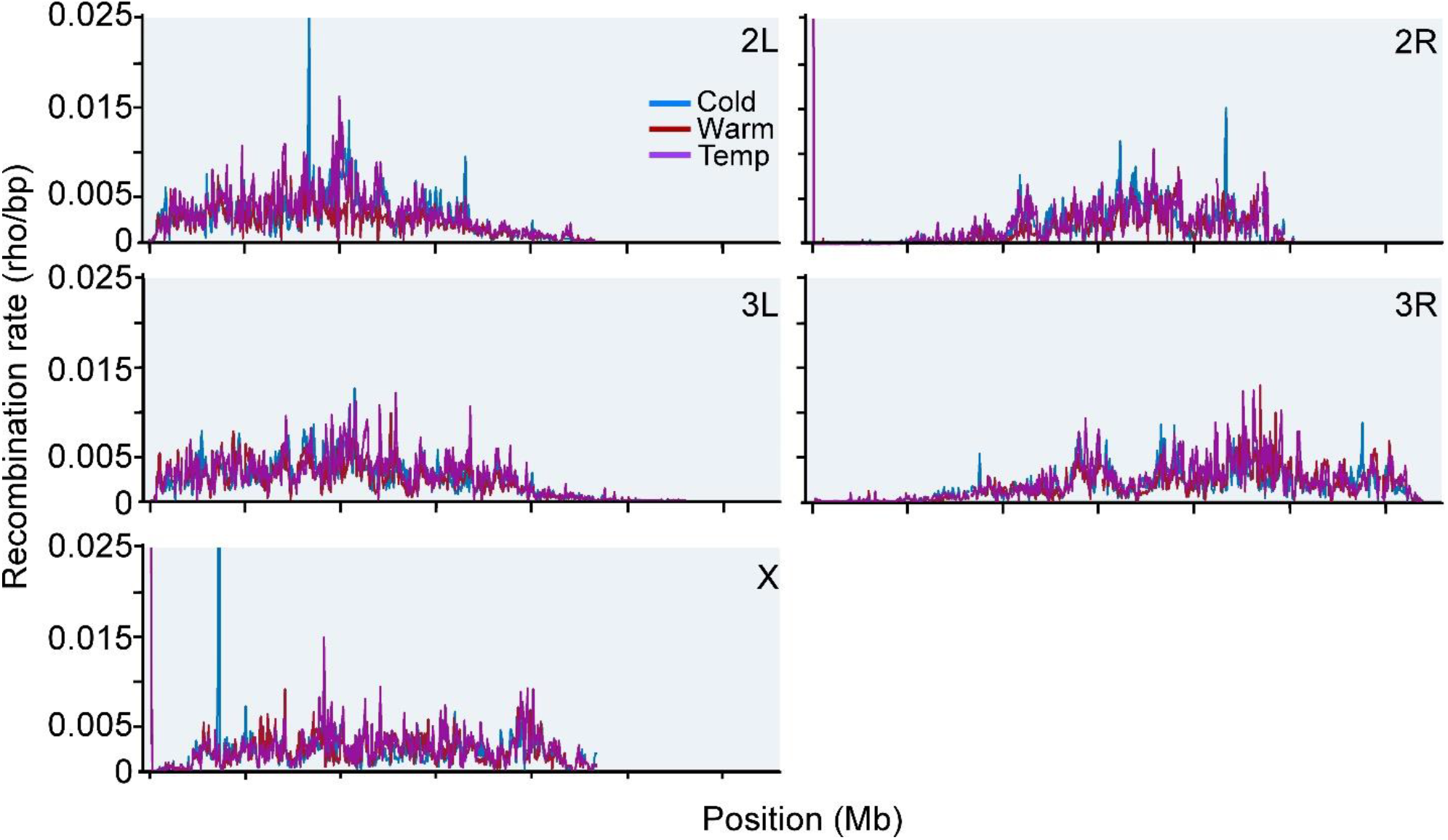
LDhelmet recombination rate estimate comparisons between the three populations (Cold, Warm, and Temp) for the five major chromosome arms averaged at 50kb intervals.

**Figure 4.**
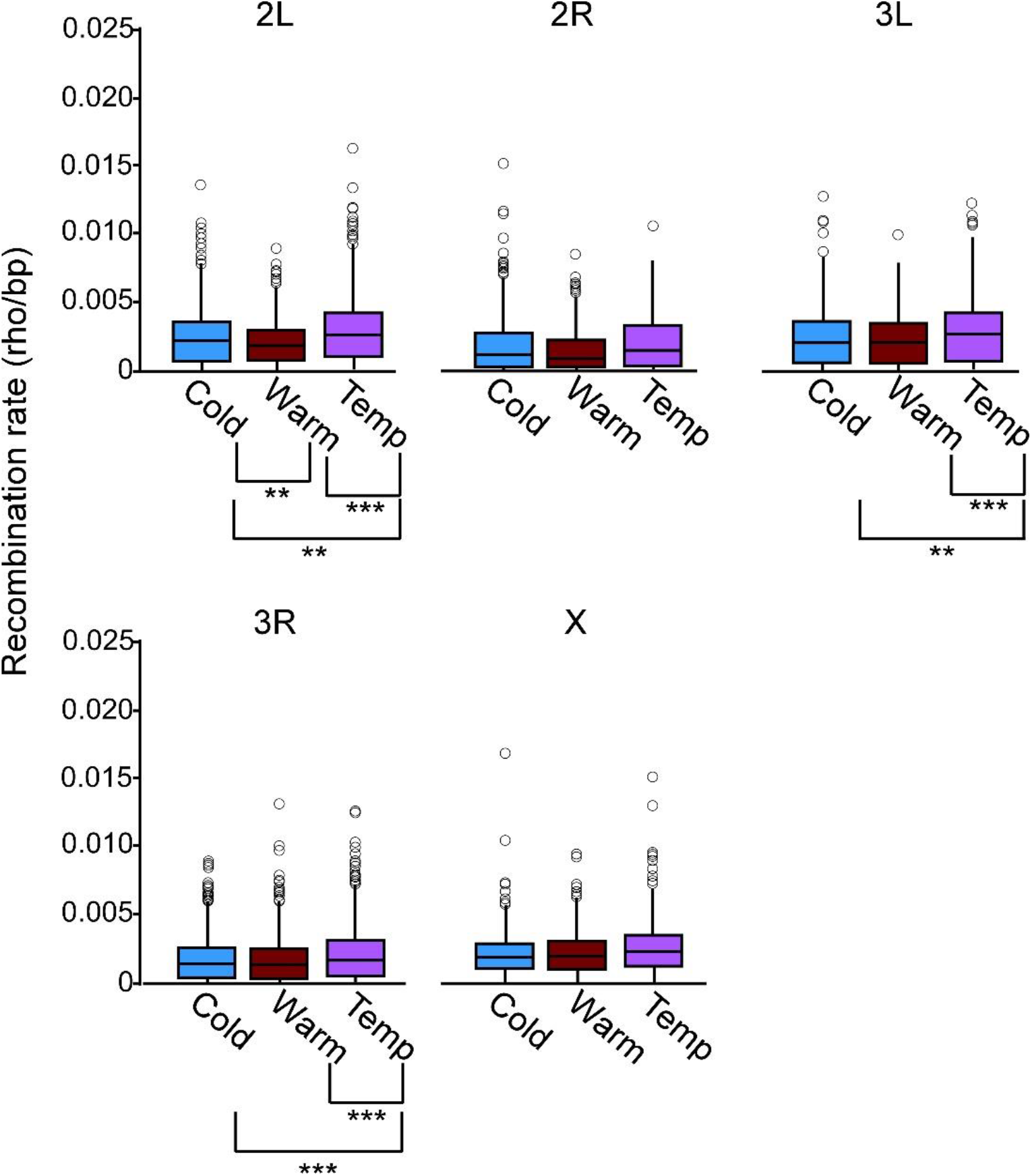
Recombination rates distribution across the five major chromosome arms averaged at 50kb intervals for the Cold, Warm and Temp populations. Most extreme outliers are omitted from figure. Population comparisons with significant differences in recombination rates are indicated with brackets. **P<0.05, ***P<0.001 (Tukey’s HSD test for all comparisons).

We next extended this analysis to broader scales to include 100kb, 200kb, and 500kb intervals (Table 3B-D, Supplemental Figure 1). In all cases the Warm vs Temp comparison continued to exhibit the highest correlations at all chromosomes with the exception of chromosome 2R at 500kb. At all intervals, the lowest correlations for all three comparisons were observed on the X chromosome and a general increase in correlation was not observed at broader scales across any chromosomes. These results suggest that recombination rate variation within chromosomes between the three populations is conserved at broader scales. At increasingly broader scales, significant differences in recombination rates on chromosome arms 2L, 2R, 3L, and 3R were observed at 100kb (P<0.01 in all instances; one-way analysis of variance), while at 200kb, significant differences are observed on chromosome arms 2L, 2R, and 3R (P<0.05 in all instances; one-way analysis of variance) and at 500kb on chromosome 2L (P<0.05; one-way analysis of variance). In most cases higher rates in the Temp population appear to be the main contributor to these differences based on post-hoc pairwise comparisons (Supplemental Figure 2A-C). Collectively these data indicate highest historical rates in the Temp population overall.

### Nucleotide Diversity Correlates

Associations between recombination rate and nucleotide diversity have previously been noted at varying levels in several species including those of *Drosophila* (Begun and Aquadro, 1992; Charlesworth and Campos, 2014; Kartje et al., 2020; Cai et al., 2009; Corbett-Detig et al., 2015) with decreases in nucleotide diversity being associated with the effects of linked selection. Similar to previous studies (Singh et al., 2013; Smukowski Heil et al., 2015), we tested whether recombination rate and nucleotide diversity were correlated in our statistical LDhelmet-based recombination data at fine scales in our three populations. At 50kb, 100kb, 200kb and 500kb intervals we observe moderate statistical correlations for all three populations for the autosomes (Table 4A-D). In the Cold and Temp populations, we observe a general but weak increase in correlation at the larger interval sizes with highest values observed at 500kb. This trend however was not as prominent in the Warm population for those chromosomes. The weakest correlations were observed in the X-chromosome with a decrease in correlation values at broader scales becoming statistically insignificant at 500kb in two of the three populations. This would suggest that there is only a weak link between diversity and recombination overall in this chromosome. Outside of this however, we see a trend in the three populations, especially the Cold and Temp in which recombination likely contributes significantly to nucleotide diversity at various scales, most notably on chromosomes 2R and 3L. Interestingly, pairwise population comparisons uncovered large areas of divergence on these two chromosomes between the three populations that appeared to overlap areas of reduced nucleotide diversity (Winbush and Singh, 2021). Therefore, we conclude a link between fine scale recombination rates and linked selection in the three populations for four of the five major chromosomes.

**Table 4.**
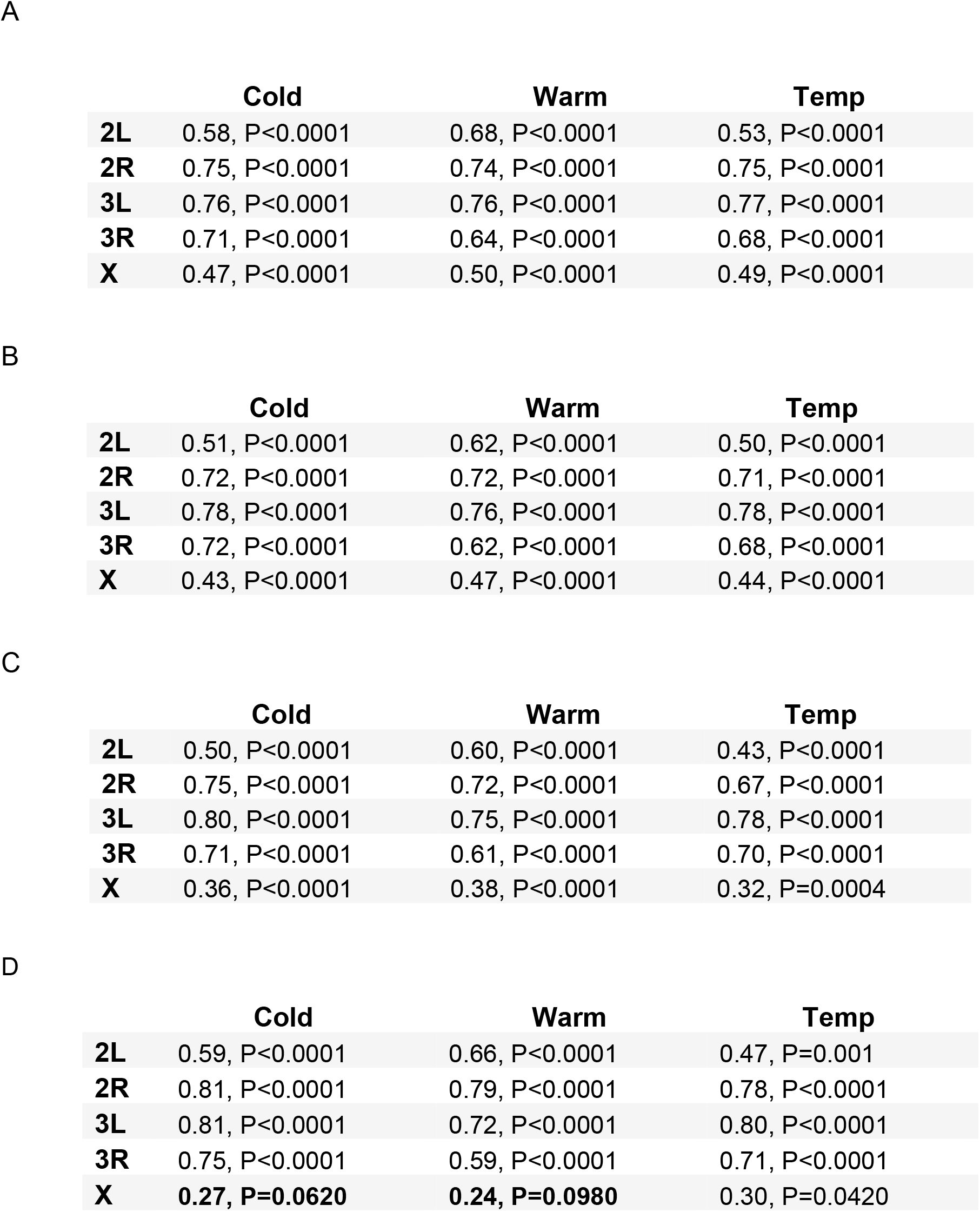
Comparison of LDhelmet-estimated recombination rates to nucleotide diversity estimates at (A) 50kb, (B) 100kb, (C) 200kb and (D) 500kb intervals for the three populations (Cold, Warm, and Temp) for the five major chromosome arms. Entries (separated by commas) as follows: Spearman’s rho, P-value. Bold entries depict entries that are not significantly correlated.

|  | Cold | Warm | Temp |
| --- | --- | --- | --- |
| 2L | 0.58, P<0.0001 | 0.68, P<0.0001 | 0.53, P<0.0001 |
| 2R | 0.75, P<0.0001 | 0.74, P<0.0001 | 0.75, P<0.0001 |
| 3L | 0.76, P<0.0001 | 0.76, P<0.0001 | 0.77, P<0.0001 |
| 3R | 0.71, P<0.0001 | 0.64, P<0.0001 | 0.68, P<0.0001 |
| X | 0.47, P<0.0001 | 0.50, P<0.0001 | 0.49, P<0.0001 |

B
|  | Cold | Warm | Temp |
| --- | --- | --- | --- |
| 2L | 0.51, P<0.0001 | 0.62, P<0.0001 | 0.50, P<0.0001 |
| 2R | 0.72, P<0.0001 | 0.72, P<0.0001 | 0.71, P<0.0001 |
| 3L | 0.78, P<0.0001 | 0.76, P<0.0001 | 0.78, P<0.0001 |
| 3R | 0.72, P<0.0001 | 0.62, P<0.0001 | 0.68, P<0.0001 |
| X | 0.43, P<0.0001 | 0.47, P<0.0001 | 0.44, P<0.0001 |

C
|  | Cold | Warm | Temp |
| --- | --- | --- | --- |
| 2L | 0.50, P<0.0001 | 0.60, P<0.0001 | 0.43, P<0.0001 |
| 2R | 0.75, P<0.0001 | 0.72, P<0.0001 | 0.67, P<0.0001 |
| 3L | 0.80, P<0.0001 | 0.75, P<0.0001 | 0.78, P<0.0001 |
| 3R | 0.71, P<0.0001 | 0.61, P<0.0001 | 0.70, P<0.0001 |
| X | 0.36, P<0.0001 | 0.38, P<0.0001 | 0.32, P=0.0004 |

Table 4. Comparison of LDhelmet-estimated recombination rates to nucleotide diversity estimates at (A) 50kb, (B) 100kb, (C) 200kb and (D) 500kb intervals for the three populations (Cold, Warm, and Temp) for the five major chromosome arms.
|  | Cold | Warm | Temp |
| --- | --- | --- | --- |
| 2L | 0.59, P<0.0001 | 0.66, P<0.0001 | 0.47, P=0.001 |
| 2R | 0.81, P<0.0001 | 0.79, P<0.0001 | 0.78, P<0.0001 |
| 3L | 0.81, P<0.0001 | 0.72, P<0.0001 | 0.80, P<0.0001 |
| 3R | 0.75, P<0.0001 | 0.59, P<0.0001 | 0.71, P<0.0001 |
| X | **0.27, P=0.0620** | **0.24, P=0.0980** | 0.30, P=0.0420 |

### Identification and mapping of Hotspots

Current evidence suggests that *Drosophila* both lack recombination hotspots on the same scale as observed in humans, and experience generally higher levels of background recombination (Manzano-Winkler et al., 2013; Comeron et al., 2012; Singh et al., 2009; Myers et al., 2005). However, regions exhibiting large scale variation in recombination rate, including areas of recombination elevated multi-fold over background have been observed in *D. melanogaster* and *D. pseudoobscura* (Comeron et al., 2012; Cirulli et al., 2006; Singh et al., 2009). At fine scales, examination of regions between 0.5 and 6.8kb in LDhelmet-estimated data identified 10 putative hotspots in the *D. melanogaster* RA and RG populations and 19 putative hotspots were identified in *D. pseudoobscura* (Chan et al., 2012; Smukowski Heil et al., 2015). We thus wanted to investigate whether we could find similar evidence of putative ‘warm-spots’ in our three populations. Searching for regions between 0.5 and 7kb in which recombination rate exceeded 10-fold the background average for that chromosome arm resulted in the identification of 9, 11, and 13 putative warm-spots in the respective Cold, Warm and Temp populations (Table 5A-C). These warm-spots included both genic and intergenic regions with no overlap observed between the three populations.

**Table 5.**
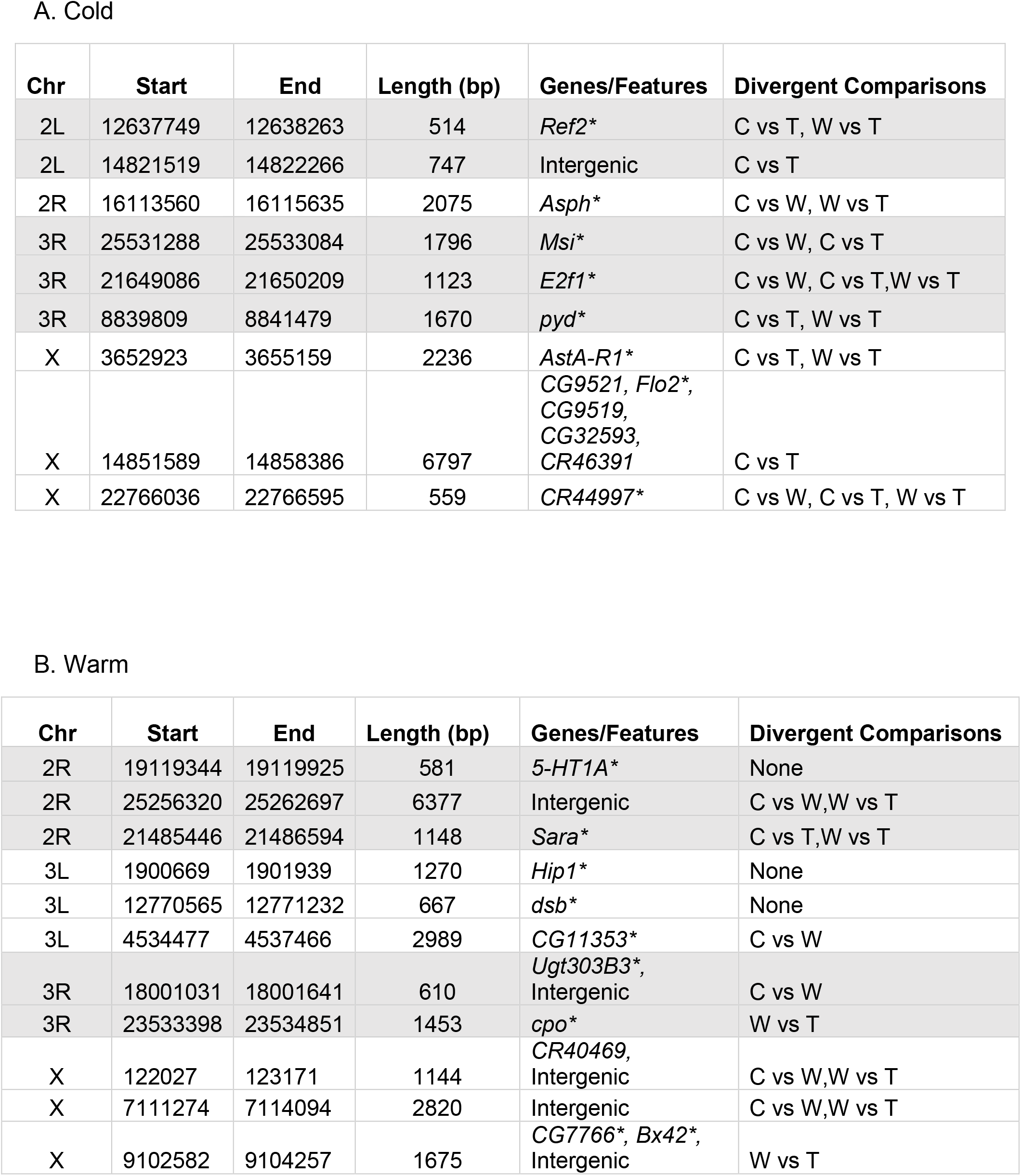
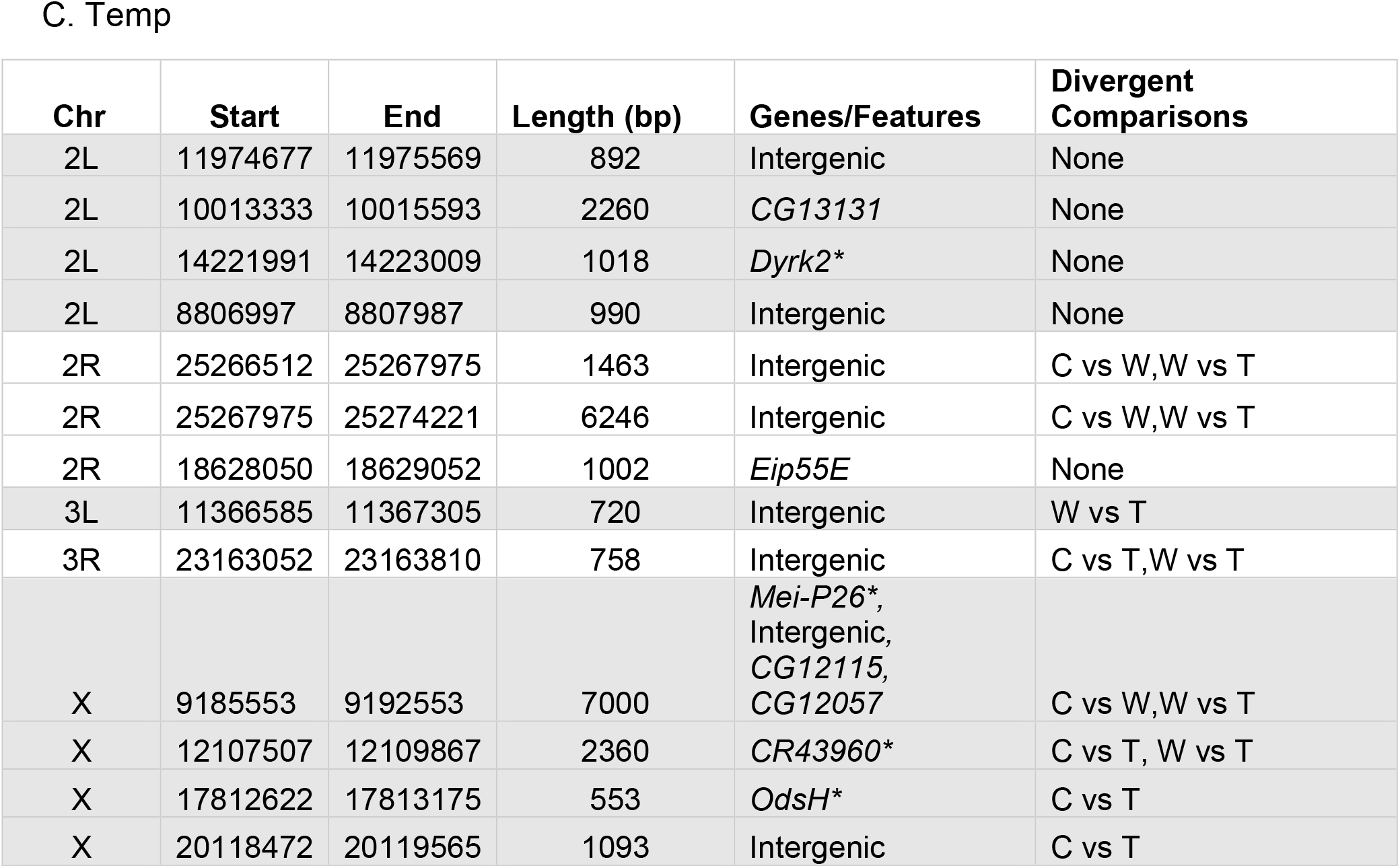
Putative warm-spots identified in the three populations (A) Cold, B (Warm) and (C) Temp based derived from LDhelmet recombination rate estimates. Columns depict the chromosome, start, end, base-pair length, and associated features of each warm-spot. Asterisks denote genes previously identified as divergent in previous study (Winbush and Singh et al., 2021). Final column depicts population comparisons for which the regions encompassing the warm-spot in question were previously identified as divergent between the populations. (C vs W: Cold vs Warm,C vs T: Cold vs Temp,W vs T: Warm vs Temp)

A. Cold
| Chr | Start | End | Length (bp) | Genes/Features | Divergent Comparisons |
| --- | --- | --- | --- | --- | --- |
| 2L | 12637749 | 12638263 | 514 | *Ref2* * | C vs T, W vs T |
| 2L | 14821519 | 14822266 | 747 | Intergenic | C vs T |
| 2R | 16113560 | 16115635 | 2075 | *Asph* * | C vs W, W vs T |
| 3R | 25531288 | 25533084 | 1796 | *Msi* * | C vs W, C vs T |
| 3R | 21649086 | 21650209 | 1123 | *E2f1* * | C vs W, C vs T, W vs T |
| 3R | 8839809 | 8841479 | 1670 | *pyd* * | C vs T, W vs T |
| X | 3652923 | 3655159 | 2236 | *AstA-R1* * | C vs T, W vs T |
| X | 14851589 | 14858386 | 6797 | *CG9521*, *Flo2* *,<br>*CG9519*,<br>*CG32593*,<br>*CR46391* | C vs T |
| X | 22766036 | 22766595 | 559 | *CR44997* * | C vs W, C vs T, W vs T |

B. Warm
| Chr | Start | End | Length (bp) | Genes/Features | Divergent Comparisons |
| --- | --- | --- | --- | --- | --- |
| 2R | 19119344 | 19119925 | 581 | *5-HT1A* * | None |
| 2R | 25256320 | 25262697 | 6377 | Intergenic | C vs W, W vs T |
| 2R | 21485446 | 21486594 | 1148 | *Sara* * | C vs T, W vs T |
| 3L | 1900669 | 1901939 | 1270 | *Hip1* * | None |
| 3L | 12770565 | 12771232 | 667 | *dsb* * | None |
| 3L | 4534477 | 4537466 | 2989 | *CG11353* * | C vs W |
| 3R | 18001031 | 18001641 | 610 | *Ugt303B3* *,<br>Intergenic | C vs W |
| 3R | 23533398 | 23534851 | 1453 | *cpo* * | W vs T |
| X | 122027 | 123171 | 1144 | *CR40469*,<br>Intergenic | C vs W, W vs T |
| X | 7111274 | 7114094 | 2820 | Intergenic | C vs W, W vs T |
| X | 9102582 | 9104257 | 1675 | *CG7766* *, *Bx42* *,<br>Intergenic | W vs T |

Table 5. Putative warm-spots identified in the three populations (A) Cold, B (Warm) and (C) Temp based derived from LDhelmet recombination rate estimates.
| Chr | Start | End | Length (bp) | Genes/Features | Divergent Comparisons |
| --- | --- | --- | --- | --- | --- |
| 2L | 11974677 | 11975569 | 892 | Intergenic | None |
| 2L | 10013333 | 10015593 | 2260 | *CG13131* | None |
| 2L | 14221991 | 14223009 | 1018 | *Dyrk2\** | None |
| 2L | 8806997 | 8807987 | 990 | Intergenic | None |
| 2R | 25266512 | 25267975 | 1463 | Intergenic | C vs W,W vs T |
| 2R | 25267975 | 25274221 | 6246 | Intergenic | C vs W,W vs T |
| 2R | 18628050 | 18629052 | 1002 | *Eip55E* | None |
| 3L | 11366585 | 11367305 | 720 | Intergenic | W vs T |
| 3R | 23163052 | 23163810 | 758 | Intergenic | C vs T,W vs T |
| X | 9185553 | 9192553 | 7000 | *Mei-P26\**,<br>Intergenic,<br>*CG12115*,<br>*CG12057* | C vs W,W vs T |
| X | 12107507 | 12109867 | 2360 | *CR43960\** | C vs T, W vs T |
| X | 17812622 | 17813175 | 553 | *OdsH\** | C vs T |
| X | 20118472 | 20119565 | 1093 | Intergenic | C vs T |

### Increased recombination rate in regions undergoing selection

We previously identified multiple chromosomal regions that were divergent in the three population comparisons (Cold vs Warm, Cold vs Temp, Warm vs Temp) using a 100kb sliding window approach that took linkage into account (Winbush and Singh, 2021). As these represent regions that were likely subject to selection during the experimental evolution period, we expected the warm-spots we identified to overlap these regions. Consistent with this expectation, the majority of warm-spots regions we identified did overlap with regions identified as divergent within at least one population comparison (Table 5A-C). This was most prominent in the Cold population warm-spots, however, the Warm and Temp populations also exhibited warm-spots meeting this criterion. This would suggest that during our experimental evolution period, certain regions subject to selection also showed enhanced recombination. It remains to be seen whether genes in these specific regions act as recombination rate modifiers however many of the genes associated with our warm-spots were previously identified as potential candidates based on differences in associated SNP allele frequencies (Winbush and Singh, 2021; Table 5A-C asterisks).

### Correcting for effects of Selection on LD-based recombination estimates

Our work on these three populations previously identified multiple regions of divergence between the populations indicating possible areas of direct or indirect selection especially in areas of reduced nucleotide diversity (Winbush and Singh, 2021). Therefore, it is possible that our estimates of recombination using LD were affected by selection in these three populations during the experimental evolution period. Previous studies for example have shown that in the context of positive selection, LD-based methods tend to slightly underestimate recombination rates (Smith and Fearnhead, 2005; Chan et al., 2012) and may lead to false inferences of warm-spots (Reed and Tishkoff, 2006). Therefore, to test for the effects of selection on our population recombination estimates we removed from our dataset all SNPs that we previously identified as divergent in any of the three population comparisons (Winbush and Singh 2021) then repeated our LDhelmet analysis, focusing on any differences in effects the new dataset might have on estimates of recombination rate variation and putative hotspot identification. Fine scale recombination maps generated with these changes looked subjectively similar to those of our original with the most notable changes being the loss of several peaks/outliers (Compare Figure 1 to Supplemental Figure 3). For example, a prominent peak on the X chromosome at approximately 4 MB in the Cold population at all scales is noticeably absent (compare Supplemental Figures 1 and 4). An examination of recombination rate variation between populations revealed similar trends to those previously observed at fine (Figure 5), and broader scales (Supplemental Figure 4A-C). Correlation analysis revealed similar trends with recombination rates being highly correlated between all three populations at the scales analyzed (50kb, 100kb, 200kb, 500kb). However, the higher correlations were observed between the Warm vs Temp and Cold vs Temp population comparisons (Supplemental Table 1). Like previous results, correlations were lowest on the X-chromosome for all three comparisons.

**Figure 5.**
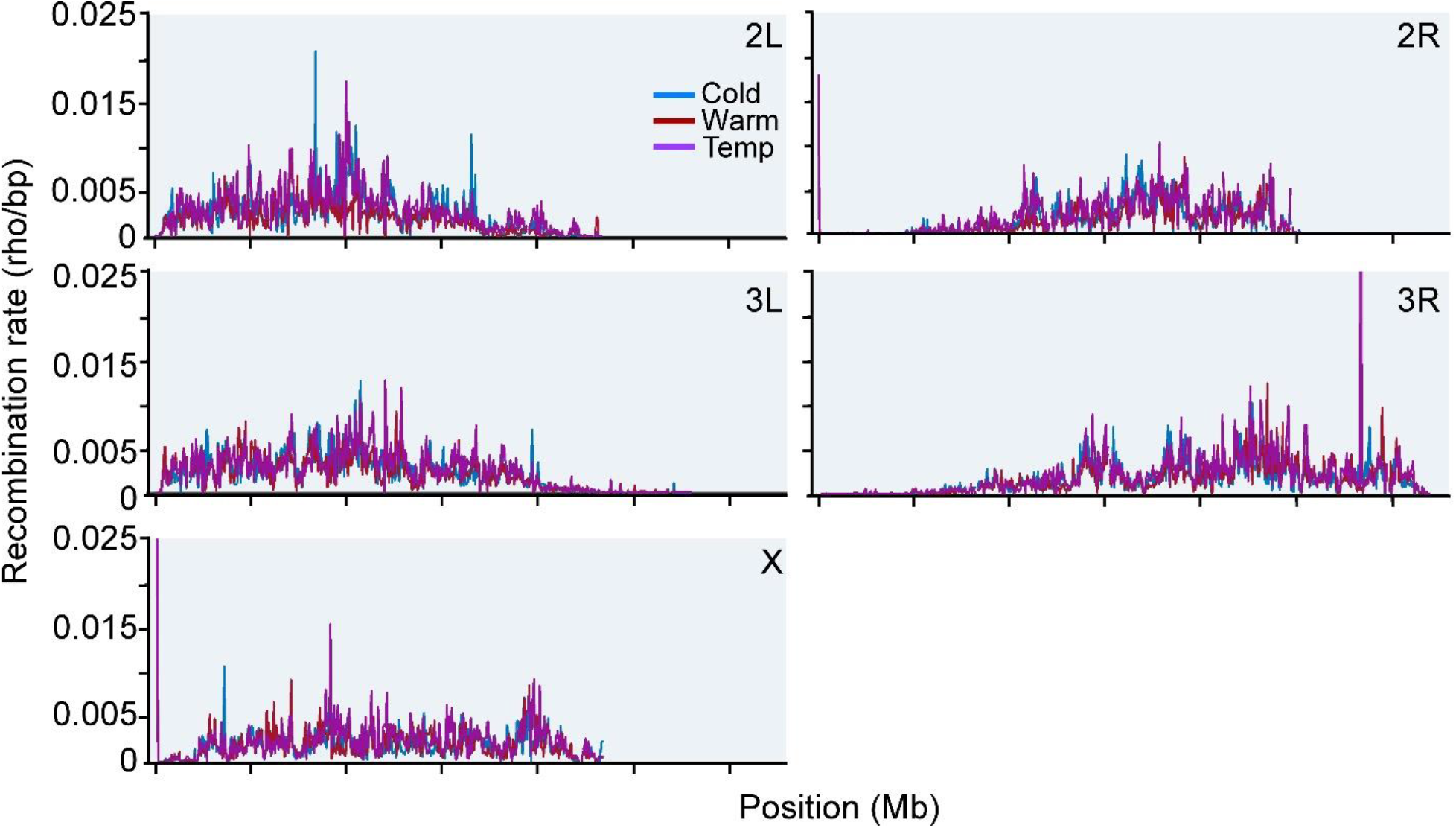
LDhelmet recombination rate estimate comparisons between the three populations (Cold, Warm, and Temp) for the five major chromosome arms averaged at 50kb intervals performed on genome sequences for which SNPs previously identified as divergent when comparing any of the three populations (Cold vs Warm, Cold vs Temp, Warm vs Temp) have been removed.

We further assessed the three populations to see if the previously observed differences in recombination rate persisted following removal of the divergent SNPs. At 50kb scale, we observed significant differences in recombination rate between the three populations for all autosomes but not X (P<0.001 in all instances; one-way analysis of variance). In addition, post hoc pairwise comparisons revealed significant differences for each population comparison for chromosome arms 2L and 2R and two out of three comparisons for chromosome arms 3L and 3R with higher rates in the Temp population appearing to be the main cause (Supplemental Figure 5A). At broader scales (Supplemental Figure 5B-D) we observed a similar trend to the earlier data. At 100kb, the significant differences in recombination rate were observed once again for all chromosomes except for X (P<0.05 in all instances; one-way analysis of variance), with post-hoc pairwise comparisons again indicating higher rates in the Temp population being the main cause. As expected, at broader scales of 200kb and 500kb, recombination rates were less divergent with only chromosomes 2L, 2R, and 3L showing significant differences at 200kb and chromosome 2L at 500kb (P<0.05 in all instances; one-way analysis variance) with the Temp population showing higher rates (Supplemental Figure 5B-D).

We also examined whether the warm-spots we previously identified persisted or changed in our modified dataset. Using the same methods, we identified 6, 12, and 14 putative warm-spots respectively in the Cold, Warm and Temp populations (Supplemental Table 2). Compared to previous results the most notable change is the decrease in putative warm-spots in the Cold population along with changes in the locations of warm-spots and associated genes. In several cases however, there was enough overlap between warm-spot regions identified in our earlier dataset that resulted in the same genes being annotated (Compare Table 5 and Supplemental Table 2 red highlights). Furthermore, as in the previous dataset, the majority of putative warm-spot regions overlapped regions previously identified as divergent and under possible selection.

In summary, despite removal the divergent SNPs in question, observed patterns and variations in recombination were largely similar with the biggest effect being in the identified putative warm-spots. This would seem to verify studies indicating the limitations of using LD-based methods to identify warm-spots in regions where selection may be occurring. As such, independent verification of these regions using empirical methods is warranted. Overall, however, our data support the general observation that recombination rate distribution is conserved during the selective period at broad scales, and more importantly areas under selection are subjected to enhanced recombination rates at fine scales.

## Discussion

The purpose of this study was to test whether regions undergoing selection to a new environment also show increases in recombination. Using LD-based methods to assess historical recombination rates coupled with our experimental evolution assay, our study has demonstrated how fine scale recombination landscapes can change in response to selection. The use of statistical based LD methods is advantageous over empirical methods in terms of time and cost; however, it must be noted that these methods represent indirect estimates of recombination. As noted earlier and in previous studies, patterns of LD are also influenced by other factors such as genetic drift, demographic history and/or bottlenecks, and selection. In our three experimental populations, genetic drift is less likely to be a factor due to the level at which we assessed fine-scale recombination rate using a pooled population approach. However, the effects of both selection and demography are potential factors.

In the case of selection, we note that LD-based methods including LDhelmet are still robust to the effects of selection with, at worse, a tendency for such methods to slightly underestimate recombination rates in these scenarios (Smith and Fearnhead, 2005; Chan et al., 2012). Nevertheless, we still tested for the effects by selection by re-running LDhelmet on our data following removal of SNPs that were previously shown to be divergent in any of our three pairwise population comparisons. Overall, the same patterns of fine scale recombination rate variation were observed among the three populations. The most notable differences concern the identification of putative recombination warm-spots in our populations. Although overlap to earlier results with these SNPs retained is noted in many of the warm-spots identified, removal of divergent SNPs did affect the number and location of putative warm-spots identified. Most notably we note fewer warm-spots identified in Cold population and differences in warm-spot location and number in the Warm and Temp populations. Despite this several regions and associated genomic features are conserved. Whether these differences indicate that our experimental design and results are indeed affected by selection cannot be determined. For instance, extensive simulations of LDhelmet demonstrate that warm-spot detection requires at least a few SNPs to be present within and near the edges of putative warm-spot regions (Chan et al., 2012). In this case removal of multiple SNPs is likely to affect the location and distribution of warm-spots in general compared to data in which those SNPs are left intact. This coupled with extensive data showing that selection likely results in an underestimation in warm-spot detection lends confidence to our results. Furthermore, the general observation that the three populations vary in warm-spots location and number whether the SNPs in question are left intact or removed supports our conclusions that the three populations have diverged in their fine scale recombination maps in response to selection.

Concerning demography, we note that our experimental evolution experiment spans a comparatively short time course of only three years compared to other studies utilizing LDhelmet that compare and infer historical recombination rate differences at the species level. Also, there is no sequencing data from the progenitor population used to initiate our three populations from which comparisons can be made. However, given that LDhelmet is still robust in the context of recent selective sweeps we remain confident that our data represent accurate estimations of the historical recombination rates during that period. Most importantly however, we note that our estimates and the empirical datasets are highly correlated at the scales tested for the five major chromosomes with correlation values slightly higher at the broader scale (Table 1), with the greatest disparity noted on the X chromosome due to the absence of several peaks in our dataset compared to that of the empirical and contributing to variation observed in the empirical dataset. We conclude therefore that we have an accurate representation of fine-scale recombination rate variation for our three populations.

### Variation in recombination rate between our three populations

The conclusion that our three populations diverged in recombination at fine scales in response to selection is most evident by our observations that at fine scales we see the greatest variation between the three populations with higher overall average rates in the Temp population (Figure 4; Supplemental Figure 2). This was most noticeable on chromosomes 2L, 3L and 3R and disappeared at larger scales. Interestingly however, all three populations are robustly correlated at all scales (Table 2). Slightly weaker correlation values were observed in pairwise comparisons involving the Cold population and lower values were likewise observed in all three pairwise comparisons in the X chromosome. From this data we conclude that at fine scales, the Temp population exhibits higher average recombination rates with statistical significance noted at finer scales for chromosomes 2L, 3L, and 3R while the Cold population exhibits slightly greater differences in terms of variation across the genome at fine scales.

Interestingly, the increased recombination rates observed for the Temp population are consistent with the hypothesized changes in recombination rate for this population resulting from the fluctuating temperature regime (Kohl and Singh, 2018). In fluctuating environments, allelic combinations may be beneficial in one environment and deleterious in the future environment. If this occurs in a cyclical environment with sufficiently long periodicity and recombination rate is condition-dependent, it was thought that higher recombination rates would evolve in populations subjected to the fluctuating environment (Kohl and Singh, 2018). Although this was not borne out when examining recombination rates within the 20.4cM region of chromosome 3R between the *ebony* and *rough* markers (Kohl and Singh. 2018) we do observe this in our fine scale data. Overall, our data supports the observation that the fine scale recombination landscapes have diverged in the three populations in response to selection in contrast to broadscale recombination rates.

### Fine scale recombination rates and selection

An overarching theme in this and other studies concerns the existence of recombination rate modifiers in *Drosophila* which become favored in response evolutionary change. Temperature and stress represent strong selective pressures, and both have been shown to increase recombination rate in *Drosophila* (Plough, 1917; Parsons, 2008; Bomblies et al., 2015). Indirect selection for increased recombination in response to selection due to these pressures may be favored in conditions of weak epistasis (Otto and Lenormand, 2002). In support of this we note an overlap between many of our putative warm-spots and regions previously shown to be divergent in the three pairwise population comparisons suggesting enrichment of recombination in areas subject to selection in response to the temperature regimes. This trend is apparent in all three populations despite variation in warm-spots and location between the three populations. Furthermore, a substantial number of genes associated with our warm-spots were previously associated with SNPs with divergent allele frequencies in the three pairwise comparisons (Winbush and Singh 2021). However, at broader scales recombination rates were observed to be correlated with nucleotide diversity supporting the observation that at broad scales, low recombination rate is favored in areas undergoing background selection for polymorphisms with the effects of linked selection contributing to reductions in nucleotide diversity surrounding those areas.

In conclusion our data provides additional evidence addressing the overarching question of how fine scale recombination landscape are affected in response to artificial selection. In our case divergence of fine scale recombination rates is noted in response to selection with areas undergoing selection exhibiting increased recombination, while conservation at broad scales is maintained.

## Data Availability

Population-specific consensus FASTA haplotype files are available at https://github.com/ariw237/temperature_recombination

## Acknowledgements

The authors wish to thank the Singh lab group for their helpful comments on the manuscript.

