## Supplemental Figures for "Variation in fine scale recombination rate in temperature-evolved *Drosophila melanogaster* populations in response to selection"

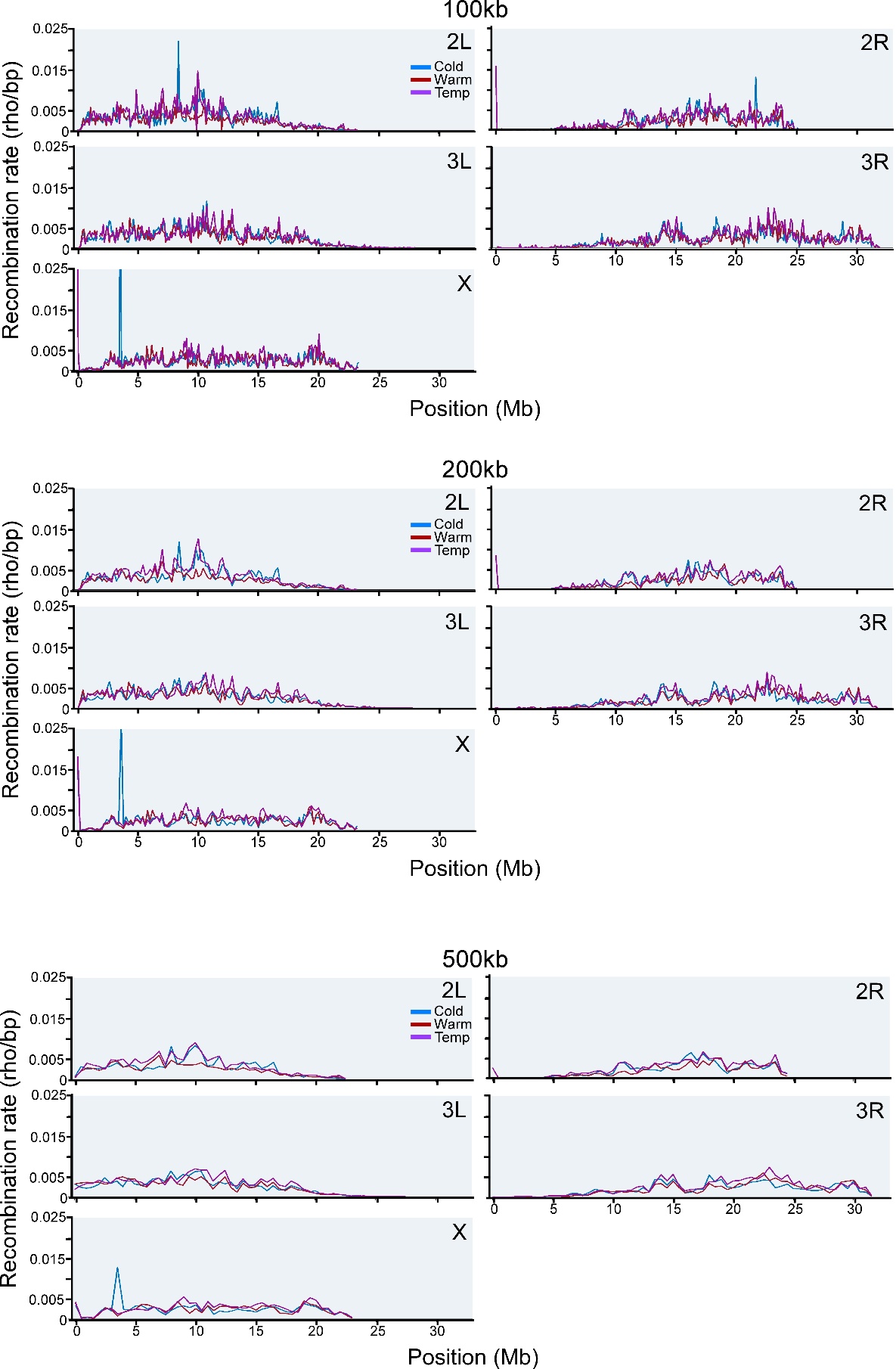

Supplemental Figure 1. LDhelmet recombination rate estimate comparisons between the three populations (Cold, Warm, and Temp) for the five major chromosome arms averaged at broader scale 100kb, 200kb and 500kb intervals.

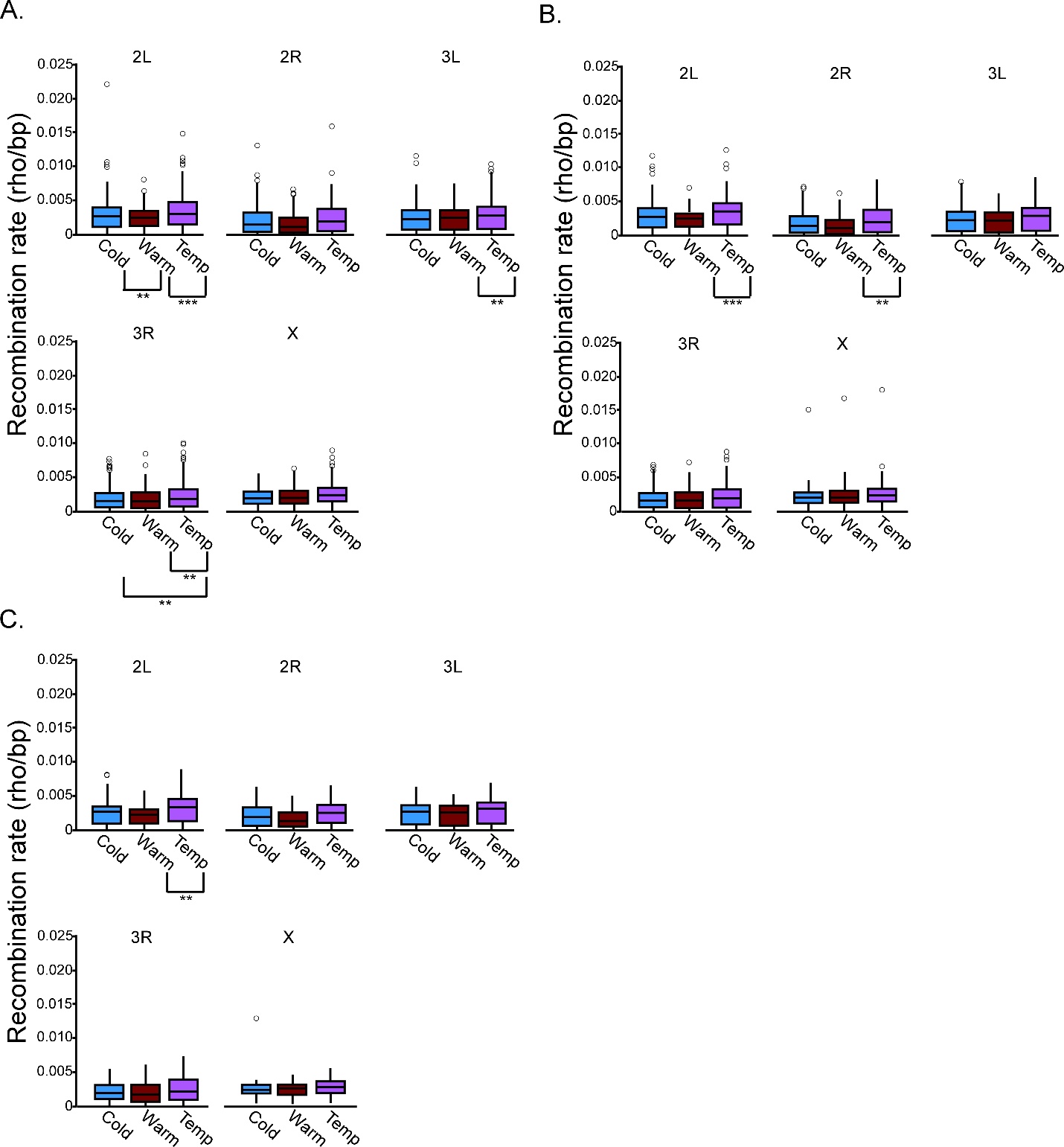

Supplemental Figure 2. Recombination rates distribution across the five major chromosome arms averaged at (A) 100kb, (B) 200kb, and (C) 500kb intervals for the Cold, Warm and Temp populations. Most extreme outliers not shown. Population comparisons with significant differences in recombination rates are indicates with brackets. **P<0.05, ***P<0.001 (Tukey’s HSD test for all comparisons).

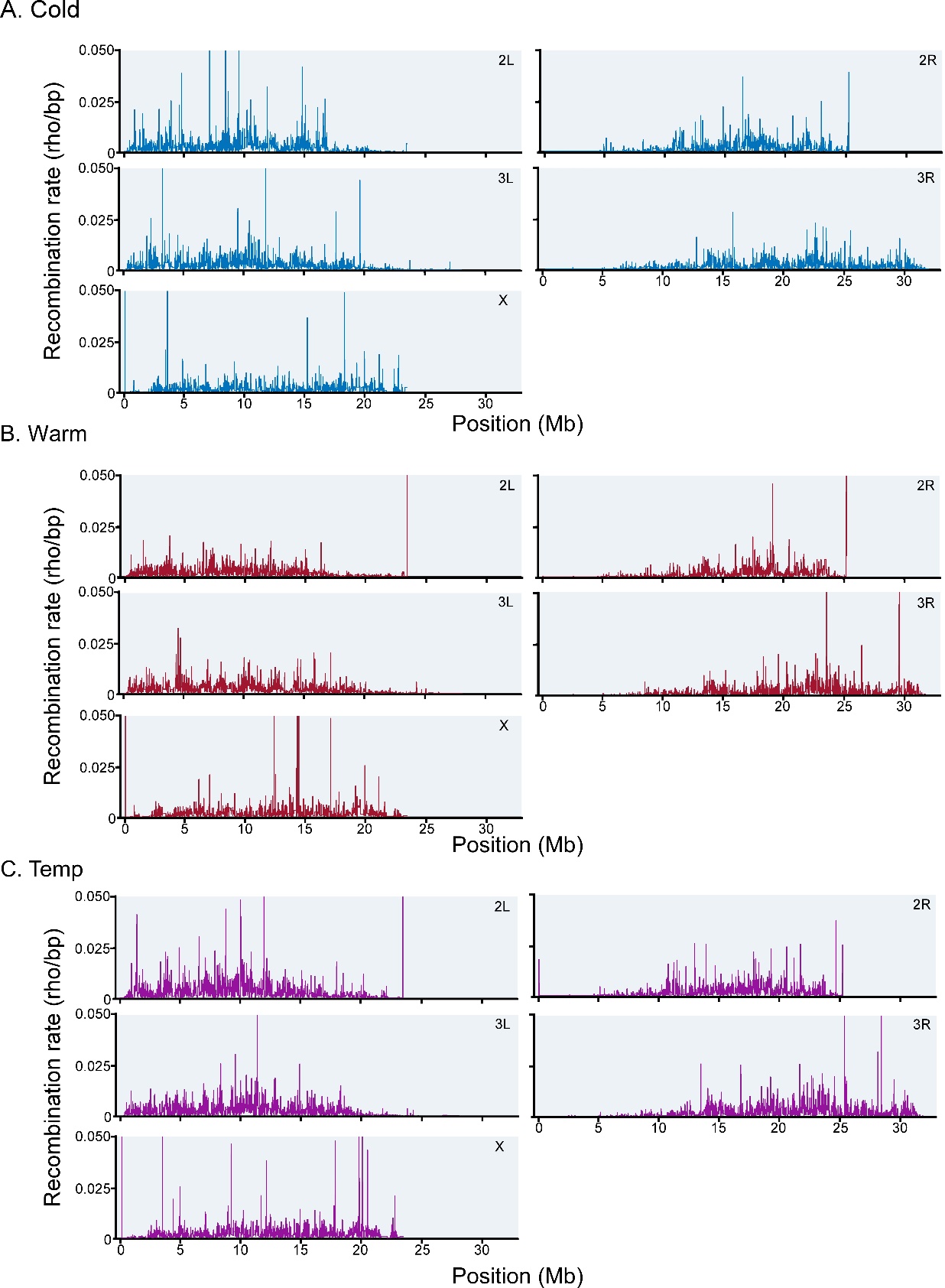

Supplemental Figure 3. LDhelmet’s fine scale recombination estimate maps performed on genome sequences for which SNPs previously identified as divergent in any of the three pairwise population comparisons (Cold vs Warm, Cold vs Temp, Warm vs Temp) have been removed. (A) Cold population. (B) Warm population. (C) Temp population.

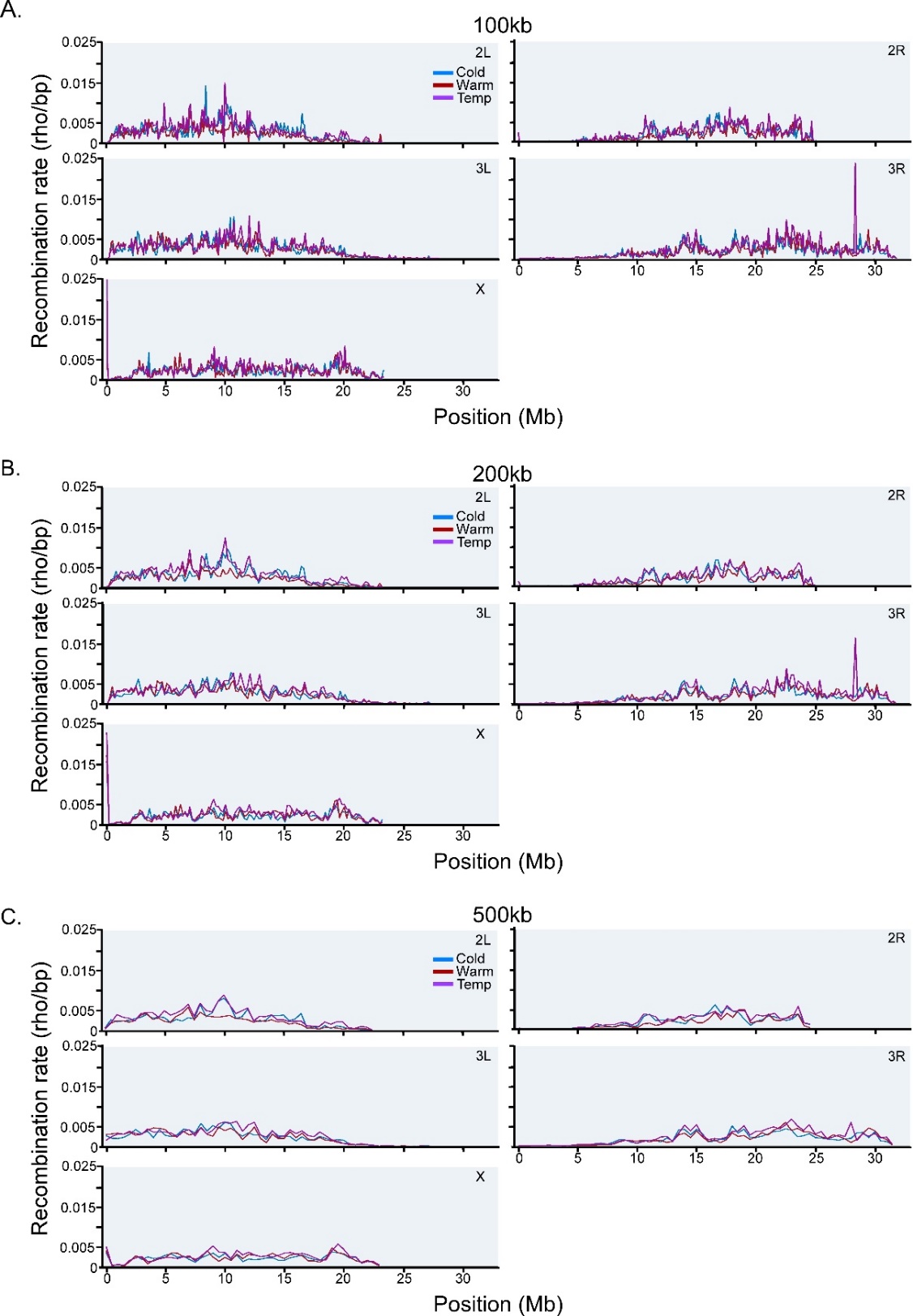

Supplemental Figure 4. LDhelmet recombination rate estimate comparisons between the three populations (Cold, Warm, and Temp) for the five major chromosome arms performed on genome sequences for which SNPs previously identified as divergent in any of the three pairwise population comparisons (Cold vs Warm, Cold vs Temp, Warm vs Temp) have been removed averaged at broader scale 100kb, 200kb and 500kb intervals.

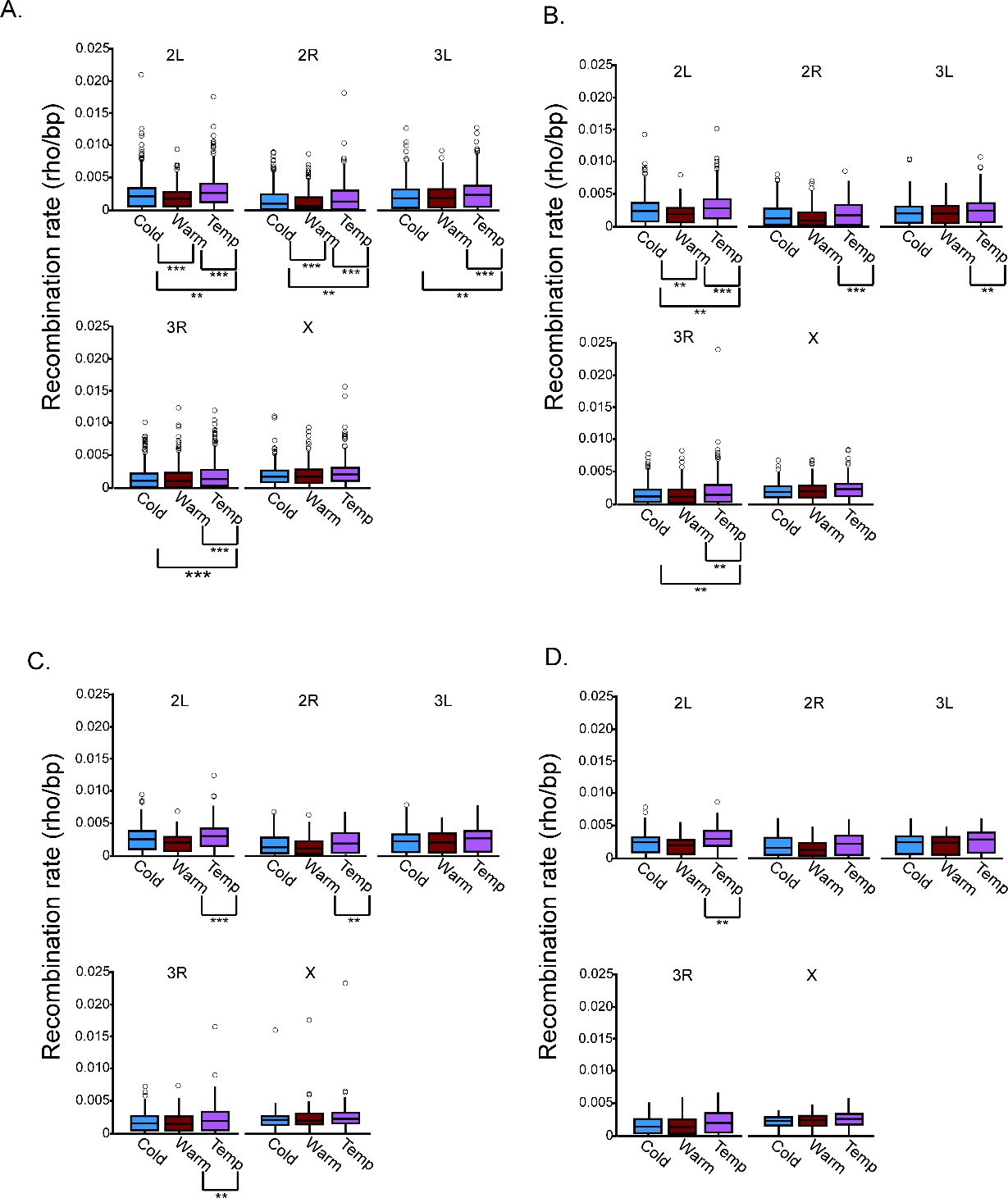

Supplemental Figure 5. Recombination rate distribution across the five major chromosome arms averaged at (A) 50kb, (B) 100kb, and (C) 200kb and (D) 500kb intervals for the Cold, Warm and Temp populations derived from LDhelmet data from genome sequences for which SNPs previously identified as divergent in any of the three pairwise population comparisons (Cold vs Warm, Cold vs Temp, Warm vs Temp) have been removed. Most extreme outliers not shown. Population comparisons with significant differences in recombination rates are indicates with brackets. **P<0.05, ***P<0.001 (Tukey’s HSD test for all comparisons).
