## Supplemental Tables for "Variation in fine scale recombination rate in temperature-evolved *Drosophila melanogaster* populations in response to selection"

A.

|  | Cold-Warm | Cold-Temp | Warm-Temp |
| --- | --- | --- | --- |
| 2L | 0.80; P<0.0001 | 0.82; P<0.0001 | 0.84; P<0.0001 |
|  | Cold: 265; Warm: 295 | Cold: 265; Temp 320 | Warm: 295; Temp: 320 |
| 2R | 0.88; P<0.0001 | 0.89; P<0.0001 | 0.90; P<0.0001 |
|  | Cold: 179; Warm: 182 | Cold: 179; Temp: 204 | Warm: 182; Temp:204 |
| 3L | 0.85; P<0.0001 | 0.86; P<0.0001 | 0.89; P<0.0001 |
|  | Cold: 248; Warm: 259 | Cold: 248; Temp: 281 | Warm: 259; Temp: 281 |
| 3R | 0.86; P<0.0001 | 0.89; P<0.0001 | 0.90; P<0.0001 |
|  | Cold: 202; Warm: 209 | Cold: 202; Temp: 227 | Warm:209; Temp 227 |
| X | 0.72; P<0.0001 | 0.72; P<0.0001 | 0.76; P<0.0001 |
|  | Cold: 172; Warm:180 | Cold:172; Temp: 194 | Warm: 180; Temp: 194 |

B.

|  | Cold-Warm | Cold-Temp | Warm-Temp |
| --- | --- | --- | --- |
| 2L | 0.80; P<0.0001 | 0.82; P<0.0001 | 0.88; P<0.0001 |
|  | Cold: 532; Warm: 592 | Cold: 532; Temp: 642 | Warm: 592; Temp: 642 |
| 2R | 0.89; P<0.0001 | 0.89; P<0.0001 | 0.90; P<0.0001 |
|  | Cold:359; Warm: 365 | Cold: 359; Temp: 407 | Warm: 365; Temp: 407 |
| 3L | 0.84; P<0.0001 | 0.85; P<0.0001 | 0.88; P<0.0001 |
|  | Cold: 497; Warm: 518 | Cold: 497; Temp: 563 | Warm: 518; Temp: 563 |
| 3R | 0.88; P<0.0001 | 0.89; P<0.0001 | 0.90; P<0.0001 |
|  | Cold: 406; Warm: 418 | Cold: 406; Temp; 455 | Warm: 418; Temp: 455 |
| X | 0.73; P<0.0001 | 0.73; P<0.0001 | 0.74; P<0.0001 |
|  | Cold: 344; Warm: 360 | Cold: 344; Temp 388 | Warm: 360; Temp: 388 |

C.

|  | Cold-Warm | Cold-Temp | | Warm-Temp |
| --- | --- | --- | --- | --- |
| 2L | 0.79; P<0.0001 | | 0.83; P<0.0001 | 0.89; P<0.0001 |
|  | Cold: 1064; Warm: 1184 | | Cold: 1064; Temp: 1285 | Warm: 1184; Temp: 1285 |
| 2R | 0.89; P<0.0001 | | 0.92; P<0.0001 | 0.91; P<0.0001 |
|  | Cold: 718; Warm: 730 | | Cold: 718; Temp: 815 | Warm: 730; Temp: 815 |
| 3L | 0.84; P<0.0001 | | 0.85; P<0.0001 | 0.87; P<0.0001 |
|  | Cold: 998; Warm: 1041 | | Cold: 998; Temp: 1131 | Warm: 1041; Temp: 1131 |
| 3R | 0.88; P<0.0001 | | 0.91; P<0.0001 | 0.89; P<0.0001 |
|  | Cold: 812; Warm: 836 | | Cold: 812; Temp: 911 | Warm: 836; Temp: 911 |
| X | 0.70; P<0.0001 | | 0.70; P<0.0001 | 0.72; P<0.0001 |
|  | Cold: 691; Warm: 724 | | Cold: 691; Temp 779 | Warm: 724; Temp:779 |

D.

|  | Cold-Warm | Cold-Temp | Warm-Temp |
| --- | --- | --- | --- |
| 2L | 0.78; P<0.0001 | 0.80; P<0.0001 | 0.92; P<0.0001 |
|  | Cold: 2703; Warm: 3006 | Cold: 2703; Temp: 3262 | Warm: 3006; Temp: 3262 |
| 2R | 0.89; P<0.0001 | 0.93; P<0.0001 | 0.92; P<0.0001 |
|  | Cold: 1788; Warm: 1819 | Cold: 1788; Temp: 2030 | Warm: 1819; Temp: 2030 |
| 3L | 0.83; P<0.0001 | 0.86; P<0.0001 | 0.90; P<0.0001 |
|  | Cold: 2497; Warm: 2603 | Cold: 2497; Temp: 2828 | Warm: 2603; Temp: 2828 |
| 3R | 0.90; P<0.0001 | 0.92; P<0.0001 | 0.89; P<0.0001 |
|  | Cold: 2030; Warm: 2091 | Cold: 2030; Temp: 2277 | Warm: 2091; Temp: 2277 |
| X | 0.70; P<0.0001 | 0.65; P<0.0001 | 0.71; P<0.0001 |
|  | Cold: 1721; Warm: 1803 | Cold: 1721; Temp: 1941 | Warm: 1803; Temp: 1941 |

Supplemental Table 1. Comparison of LDhelmet-estimated recombination rates for the three population comparisons (Cold vs Warm, Cold vs Temp, Warm vs Temp) for the five major chromosome arms at (A) 50kb interval, (B) 100kb interval, (C) 200kb, and (D) 500kb interval. Recombination rates were estimated however following removal of SNPs previously identified as divergent between any of the three populations. Entries in first row for each chromosome (separated by semicolons) as follows: Spearman’s rho, P-value while second row entries (separated by semicolons) give average number of SNPs per interval for the appropriate population.

A. Cold

| Chr | Start | End | Length | Genes/Features |  |
| --- | --- | --- | --- | --- | --- |
| 2L | 14821519 | 14822266 | 747 | Intergenic | C vs T |
| 2R | 16468170 | 16468745 | 575 | Intergenic, *CheB53b** | C vs T |
| 3L | 19673437 | 19677163 | 3726 | Intergenic | C vs T, W vs T |
| 3L | 11839037 | 11840089 | 1052 | Intergenic | None |
| X | 18317354 | 18317913 | 559 | *CG6023** | None |
| X | 15234880 | 15235883 | 1003 | Intergenic | C vs T, W vs T |

B. Warm

| Chr | Start | End | Length | Genes/Features |  |
| --- | --- | --- | --- | --- | --- |
| 2L | 23482165 | 23483533 | 1368 | Intergenic | None |
| 2L | 23483533 | 23488117 | 4584 | Intergenic | None |
| 2L | 23488117 | 23489102 | 985 | Intergenic | None |
| 2R | 19119316 | 19119925 | 609 | *5-HT1A** | None |
| 3L | 4534406 | 4537466 | 3060 | *CG11353** | C vs W |
| 3R | 23533554 | 23534851 | 1297 | *Ugt303B3*,* Intergenic | W vs T |
| 3R | 26451582 | 26454063 | 2481 | *Hex-t2,* Intergenic | W vs T |
| 3R | 29572995 | 29573557 | 562 | Intergenic | C vs T |
| X | 122027 | 123171 | 1144 | Intergenic, *CR40469* | C vs W, W vs T |
| X | 14302529 | 14303407 | 878 | Intergenic | C vs T |
| X | 17143907 | 17143907 | 821 | *RpS5a**, *CR34594..CR34597* | C vs T |
| X | 19971345 | 19974370 | 3025 | Intergenic | W vs T |

C. Temp

| Chr | Start | End | Length | Genes/Features |  |
| --- | --- | --- | --- | --- | --- |
| 2L | 10013333 | 10015565 | 2232 | *CG13131* | None |
| 2L | 10049527 | 10050135 | 608 | Intergenic, *CG33301** | None |
| 2L | 11974803 | 11975507 | 704 | Intergenic | None |
| 2L | 23487311 | 23488197 | 886 | Intergenic | None |
| 2L | 8806997 | 8807987 | 990 | Intergenic | None |
| 3L | 11366585 | 11367305 | 720 | Intergenic | C vs T |
| 3R | 28431308 | 28431914 | 606 | *Beat-VI* | C vs W, W vs T |
| X | 9189624 | 9192553 | 2929 | *CG12115*,* Intergenic, *CG12057** | C vs W, C vs T |
| X | 12107459 | 12108024 | 565 | *CR43960** | C vs T , W vs T |
| X | 17812622 | 17813175 | 553 | *OdsH** | C vs T |
| X | 20118337 | 20119565 | 1228 | Intergenic | None |
| X | 19821356 | 19824844 | 3488 | Intergenic, *SkpE** | W vs T |
| X | 12108024 | 12109867 | 1843 | *CR43960*, CG2750** | C vs T, W vs T |
| X | 19790435 | 19791277 | 842 | Intergenic | None |

Supplemental Table 2. Putative hotspots identified in the three populations (A) Cold, B (Warm) and (C) Temp based derived from LDhelmet recombination rate estimates following removal of SNPs previously identified as divergent between any of the three populations. Columns depict the chromosome, start, end, base-pair length, and associated features of each hotspot. Asterisks denote genes previously identified as divergent in previous study (Winbush and Singh et al., 2021) while genes in red text are those previously identified prior to removal of divergent SNPs (see Table 5). Final column depicts population comparisons for which the regions encompassing the hotspot in question were previously identified as divergent between the populations. (C vs W: Cold vs Warm; C vs T: Cold vs Temp; W vs T: Warm vs Temp).
